# Antarctic Teleosts With and Without Haemoglobin Behaviourally Mitigate Deleterious Effects of Acute Environmental Warming

**DOI:** 10.1101/2021.04.27.441509

**Authors:** Iskander I. Ismailov, Jordan B. Scharping, Iraida E. Andreeva, Michael J. Friedlander

**Affiliations:** Fralin Biomedical Research Institute at Virginia Tech Carilion, Roanoke, VA; Virginia Tech Carilion School of Medicine, Roanoke, VA

**Author notes:** **Co-corresponding authors:** Iskander I. Ismailov, Fralin Biomedical Research Institute at VTC, 2 Riverside Cir, Roanoke, VA 24016. Michael J. Friedlander, Fralin Biomedical Research Institute at VTC, 2 Riverside Cir, Roanoke, VA 24016.

**Keywords:** Antarctic_Notothenioids, behavioural thermoregulation, fanning, fixed action pattern, lateralization, respiratory behaviour, respiratory-locomotor coupling, startle-like behaviour

## Abstract

Recent studies forecast that many ectothermic animals, especially aquatic stenotherms, may not be able to thrive or even survive predicted climate change. These projections, however, generally do not call much attention to the role of behaviour, an essential thermoregulatory mechanism of many ectotherms. Here we characterize species-specific locomotor and respiratory responses to acute ambient warming in two highly stenothermic Antarctic Notothenioid fishes, one of which (*Chaenocephalus aceratus*) lacks haemoglobin and appears to be less tolerant to thermal stress as compared to the other (*Notothenia coriiceps*), which expresses haemoglobin. At the onset of ambient warming, both species perform distinct locomotor manoeuvres that appear to include avoidance reactions. In response to unavoidable progressive hyperthermia, fishes demonstrate a range of species-specific manoeuvres, all of which appear to provide some mitigation of the deleterious effects of obligatory thermoconformation and to compensate for increasing metabolic demand by enhancing the efficacy of branchial respiration. As temperature continues to rise, *Chaenocephalus aceratus* supplements these behaviours with intensive pectoral fin fanning which may facilitate cutaneous respiration through its scaleless integument, and *Notothenia coriiceps* manifests respiratory-locomotor coupling during repetitive startle-like manoeuvres which may further augment gill ventilation. The latter behaviours, found only in *Notothenia coriiceps*, have highly stereotyped appearance resembling Fixed Action Pattern sequences. Altogether, this behavioural flexibility could contribute to the reduction of the detrimental effects of acute thermal stress within a limited thermal range. In an ecologically relevant setting, this may enable efficient thermoregulation of fishes by habitat selection, thus facilitating their resilience in persistent environmental change.

---

There is increasing concern about the fitness of life forms on Earth in the face of rising environmental temperatures (Thomas et al., 2004; Pacifici et al., 2015). As part of that awareness, the potential vulnerability of high latitude marine ecosystems, such as the Antarctic shelf, has drawn particular attention (Clarke et al., 2012; Constable et al., 2014), as they are likely facing some of the most rapid environmental changes on the planet (Vaughan et al., 2003; Cheng et al., 2019; see, however, Turner et al., 2016). Furthermore, isolated from the rest of the world by the Antarctic Circumpolar Current, inhabitants of this ecosystem have evolved under extremely cold conditions which remained thermally stable for millions of years (Barker and Thomas, 2004). In effect, they are thought to have specialized for such a milieu by developing stenothermy, *i.e.*, traded-off the ability to adjust to even small variations in temperature (Wohlschlag, 1964; Somero and DeVries, 1967).

Based on the results of experimental studies in a variety of aquatic organisms subjected to acute warming, one perspective on the physiological mechanisms that limit adaptive capacity and thermal tolerance of ectotherms is that cardiac collapse is ultimately responsible for organismal failure at extreme temperatures (see Eliason and Anttila, 2017 for a Review). Deriving rationale from the relationship between thermal tolerance of fishes to their aerobic metabolic rates (Fry and Hart, 1948), these studies provide the basis for an *‘oxygen- and capacity-limited thermal tolerance’* theory (Pörtner, 2001) and attribute this collapse to a mismatch between the aerobic demand of the heart during thermally induced tachycardia and maximal ability to supply oxygen to the heart. Presented initially as an integrative concept comparing the effects of temperature on different organisms, this theory has lately generated discussion (Pörtner et al, 2017; Jutfelt et al., 2018) as a possible overreach attempting to merge environmentally meaningful thermal limits observed in natural habitats with acute sub-lethal warming experiments at miniscule timescales having little ecological relevance. Furthermore, experimental studies in instrumented, often sedated or restrained animals, or in perfused *in vitro* and *in situ* preparations, generally do not encompass a critical thermoregulatory mechanism of ectotherms – behaviour (Crawshaw, 1979).

In this study, we examine behavioural responses elicited by ambient warming in two highly stenothermal Antarctic teleosts, *Notothenia coriiceps* (*N. coriiceps*, Richardson 1844) and *Chaenocephalus aceratus* (*C. aceratus*, Lönnberg 1906), swimming unrestricted in a tank. These fishes belong to the related families *Nototheniidae* and *Channichthyidae*, respectively, but differ in that the blood of *C. aceratus* (also called icefish) is devoid of the oxygen transport protein haemoglobin (Ruud, 1954; Cocca et al., 1995). Consequently, haemoglobinless (Hb−) icefishes supply tissues with oxygen dissolved directly in plasma, with an estimated oxygen-carrying capacity 90% lower than the blood of haemoglobin expressing (Hb+) Antarctic fishes (Holeton, 1970). Large volumes of plasma, big hearts and elaborate vasculature (Hemmingsen, 1991) are thought to offset the Hb− state in icefishes, affording their successful habitation in the oxygen-rich Southern Ocean. Yet, compared to their Hb+ relatives, icefishes appear to be less tolerant to acute warming (Beers and Sidell, 2011), putatively due to inferior capacity to adjust cardiac performance at elevated temperatures (O’Brien et al.,2018; see, however, Joyce et al., 2018a). Hence, icefishes have been predicted to be particularly vulnerable to expected global climate change, unless behavioural thermoregulation and/or other physiological plasticity mechanisms alleviate (Huey, et al., 2012) the effects of this detriment.

Otherwise, while being considered extremely stenothermal, negatively buoyant bottom dwellers *N. coriiceps* and *C. aceratus* are both in fact eurybathic (Eastman, 2017). That is, occasionally found at depths as far down as 500 and 700 metres, respectively, both fishes prefer bathymetric ranges closer to the surface. Namely, *N. coriiceps* are most commonly found at depths less than 100 m, while *C. aceratus* are typically observed at depths between 100 and 300 m (Hureau, 1985), conceivably both performing frequent routine migrations in the water column. In all locations inhabited by both fishes from as far North as South Georgia and Bouvetøya islands (both at ~54°S) to the Antarctic Peninsula (~65.5°S) (Hureau, 1985), on the other hand, these depths correspond to a prominent thermo- and oxycline (most significant between 50 and 300 m, Fig. S1), encompassing a range of temperatures from −1.8°C to +2°C and a dissolved oxygen (DO_2_) gradient from ~10 mg·L^−1^ to ~6 mg·L^−1^. Then, the fishes must possess certain adaptive capacities that allow them to function equally well in such diverse environments. Behavioural strategies of the Hb+ and Hb− animals responding to ambient warming, however, may be different. Testing this possibility constituted the main goal of this study.

We report that both Hb+ and Hb− Antarctic fishes respond to acute ambient warming with an extensive repertoire of behaviours. Restricted in the ability to perform essential thermoregulatory behaviour such as habitat selection in the uniformly heated tank, fishes respond to progressive hyperthermia with an elaborate range of manoeuvres - relative quiescence, startle-like behaviours, aquatic surface respiration, pectoral fin fanning and pectoral fin splay, accompanied with continuous changes in ventilation. While manifestations of these behaviours are species-specific in terms of intensity, duration, thermal dependence and stereotypy, most of them are observed in both fishes. To compensate for increasing respiratory demand at the temperatures above +8°C, both fishes enhance the efficacy of their respiratory pumps. In addition, sedentary Hb− fish supplement increased ventilatory effort with intense fanning which may facilitate cutaneous respiration, whereas more agile Hb+ fish augment it with respiratory-locomotor coupling during repetitive startle-like manoeuvres. Altogether, these behavioural responses appear to constitute complementary adaptive actions to physiologically mitigate deleterious effects of obligatory thermoconformation, resulting in concerted optimization of multiple vital functions within a limited thermal range.

## MATERIALS AND METHODS

### Animals

Animals were collected during the austral fall of 2015 (April-June) off the Southwestern shore of Low Island (63°30 S, 62°42 W) and of Brabant Island in Dallmann Bay near Astrolabe Needle (64°08 S, 62°40 W). Otter trawls were used to capture both *C. aceratus* and *N. coriiceps*, and baited benthic traps - for *N. coriiceps* only, all deployed from the Antarctic Research and Supply Vessel *Laurence M. Gould* (LMG). On board of the LMG, captured animals were held segregated by the species up to 4 days in insulated 900 litre tanks (Xactics™, ON, Canada) supplied with running ambient ocean water and superfluous aeration at a rate of 17 litres per minute provided by two submersed glass-bonded silica air diffusers (Sweetwater® model AS5L, Pentair Aquatic Eco-Systems, FL, USA) and two diaphragm air pumps (24 litres per minute at 1 psi output, Sweetwater® model SL24, Pentair Aquatic Eco-Systems, FL, USA) per tank. After transfer to aquaria at the United States Antarctic Program research station, Palmer Station, fishes were kept segregated by the species in 9,000 litre tanks flowing fresh sand-filtered ocean water pumped from the Arthur Harbour at ambient temperature (−1.7°C to +1°C) for a minimum period of 72 h and up to 3 weeks prior to experiments. *N. coriiceps* were fed fish muscle blocks once every 2-3 days. Icefish do not feed in captivity (O’Brien et al., 2018), so these animals were used within two weeks of capture. All animal procedures were approved by the University of Alaska, Fairbanks Institutional Animal Care Committee (570217-9).

### Temperature ramp experiments

Behavioural experiments were performed in a custom-built 500-litre (93 cm (W) x 93 cm (L) x 74 cm (H)) flow-through experimental acrylic tank placed in a climate controlled room (with air temperature maintained between +2 °C and +4°C) and filled with 300 litres of seawater. All surfaces of the tank were additionally covered with 3.175 mm thick transparent red (#2423) acrylic (Professional Plastics, Inc., CA, USA). The rationale for the latter was based on the finding that Notothenioids lack long-wave sensitive opsin gene (Pointer et al., 2005). Therefore, used photic conditions mimic light environment corresponding to the austral mid-winter darkness in the natural habitat of the fishes, visually shield them from the experimenters, and should help to minimize stress.

Before the experiment, each specimen was allowed to acclimate overnight in the experimental tank continuously flowing with fresh ocean water (at a rate of 11-15 litres per minute) pumped from the Arthur Harbour at ambient temperature. This ensured relative consistency of thermal environment of fishes from capture to the beginning of the experiment. Baseline behaviours were recorded during this period (for at least 60 minutes just prior to warming ramp). Warming rate of ~3.2°C per hour was achieved by re-circulating tank water through the coil of a custom-made glass heat exchanger with a jacket plumbed to a heating bath-pump (AD28R-30, VWR, PA, USA) running in an external closed loop mode. During the temperature ramp, before entering the heat exchanger, re-circulated water was aerated at a rate of 17 litres per minute using two submersed glass-bonded silica air diffusers (Sweetwater® model AS5L, Pentair Aquatic Eco-Systems, FL, USA) and a diaphragm air pump (24 litres per minute at 1 psi output, Sweetwater® model SL24, Pentair Aquatic Eco-Systems, FL, USA).

Digital video images were acquired with Ethovision XT10 tracking software (Noldus Information Technologies, Inc., Netherlands) at a standard frame rate of 30 Hz using a Basler acA 1300-60gm area scan GigE camera (Basler AG, Germany) equipped with a Computar H2Z0414C-MP CCTV lens (CBC Group, NC, USA). The tank contained no gravel or any other substrate, and imaging was performed in a transparency mode (ventral view), in a 34 x34 acrylic mirror (Professional Plastics, Inc., CA, USA) placed at a shallow angle (to minimize distortions) 75 cm below the tank. Illumination was achieved with four 12 x12 heat- and flicker-free (200W HMI light output) LED flood light panels (model 1×1LS Litepanels, Vitec Group, UK) positioned 50 cm above the tank and waterproofed by lamination from both sides 36% transmission value diffusion filter (# 216, LEE Filters, UK) placed directly over the tank. Water temperature and DO_2_ were recorded synchronously using an Orion Versa Star meter and DO_2_ probe (Thermo Scientific, MA, USA). Warming and video recordings continued until loss of equilibrium (LOE) was observed. All experiments were terminal, and each animal was used in a single behavioural experiment without repetition.

### Analyses of locomotor activity

Locomotion was analysed *post hoc* in video recordings acquired in the field using an automated tracking algorithm of EthoVision XT software. Instantaneous velocities (expressed in body lengths per second, BL·s^−1^) were calculated by the program from instantaneous (*i.e.*, measured between every two consecutive frames taken 33 ms apart) displacement of the centre of the mass of the fish (determined using a proprietary algorithm of the EthoVision software). The average body lengths were 36.91 ± 1.26 cm and 47.36 ± 3.22 cm, for five *N. coriiceps* and five *C. aceratus*, respectively (mean ± SEM). Elongation ratios of fishes used in our experiments were determined as the ratio of length over width (Ward and Azizi, 2004), and constituted 8.38 ± 0.24 (mean ± SEM; n = 5) for *C. aceratus* and 4.73 ± 0.11 (mean ± SEM; n = 5) for *N. coriiceps*.

### Analyses of respiratory behaviours

Opercular movements were analysed *post hoc* by manual frame-by-frame measurements of fish head width in Adobe Premiere Pro v.5.5.2 (Adobe Inc., CA, USA) projecting field video recordings on the screen of a Dell U2410 monitor (1920 x 1200 resolution) at 3.2x magnification. All metrics were determined unremitted throughout entire experiment, except the moments when high linear or angular velocity of fish movement precluded reliable measurements. Ventilation frequencies were determined by tallying of opercular movement cycles per minute, and compared for consistency with records of counts taken at 10-15 minute time intervals during actual field experiments at Palmer Station. Opercula opening amplitude (OA), a direct proxy metric of ventilatory stroke volume, was measured as the difference in width of fish head with opercula maximally open and maximally closed, using an Apollo VCG7070 transparency film (Acco Brands, IL, USA) with a 1 mm grid. Accuracy of head width change measurements was arbitrarily set at 0.5 mm. Opercula opening time (OT), an inverse proxy metric of branchial pump suction, was calculated from the number of frames taken during transition of opercula from closed to maximally open state. Opercula opening velocity (OV) calculated as OV = OA/OT. This ratio is considered as “unidimensional” proxy metric of branchial pump velocity to characterize the efficacy of ventilation. Due to suboptimal contrast in some recordings, reliable characterization of respiration metrics was possible only in three specimens of each species. Numbers and durations of surfacing events were manually tallied *post hoc* on the frame-by-frame basis in Adobe Premiere Pro.

### Kinematics of startle-like behaviours

Spontaneous startle-like behaviours were analysed *post hoc* manually in Adobe Premiere Pro, on a frame-by-frame basis, using conventions comparable to those described above for analyses of respiratory behaviours and Apollo VCG7070 transparency film (without a grid) placed directly on a Dell U2410 monitor projecting field video recordings. Angular velocities were determined by measuring with a transparent protractor angles between vectors drawn from the centre of mass (vertex of the angle, determined using a proprietary algorithm of the EthoVision software) of the fishes to the snout at appropriate temporal resolution. For fast S-bend startle-like manoeuvres of *N. coriiceps*, a 33 ms^−1^ frame rate may result in underestimation of angular velocity.

Analyses of axial movements during thermally induced startle-like responses were done using manual sketches of the body shapes of the fishes made on transparency film on a frame-by-frame basis. For presentation purposes, ventral outlines of the turning fish were made over scanned and imported into CorelDraw (Corel Corporation, ON, Canada) images.

### Lateralization of C-bend manoeuvres in response to ambient warming

For laterality analyses, quantities and directions of C-bend manoeuvres were tallied for every 0.5°C increment of the temperature ramp in each of five experiments with Hb+ and Hb− fishes. Since tank walls can affect lateralization (Eaton and Emberly, 1991), turns that occurred near the walls were not tallied. Relative Lateralization indices (LR, %, Bisazza et al., 1998) were calculated using following equation:

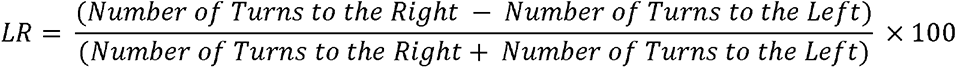

We note, however, that unlike in detour paradigm of Bisazza et al. analysing single instance of behaviour of each fish in multiple experimental trials, we analysed multiple manoeuvres of each specimen in a single continuous warming experiment within the range of temperatures in which the fish demonstrated spontaneous, repetitive startle-like behaviours. Mean LR (± SEM) was used to assess turning preference (i.e. bias in left or right turns) for 5 specimens of each species. The LR index at the level of individuals allowed to classify specimens between the extreme values of “100” (when fish turned right in all cases recorded within temperature interval corresponding to 5% T_LOE_) and “−100” (when fish turned left on all cases recorded within temperature interval corresponding to 5% T_LOE_). A mean LR near zero indicates that a given species is neither left nor right biased in its tendency to turn in a given temperature range.

### Analyses of fin movements

Duration of fanning bouts, fanning frequency (as a number of pectoral fin beats per second within a bout) and duration of pectoral fin splay were manually tallied *post hoc* on the frame-by-frame basis in Adobe Premiere Pro.

### Quantification, presentation of grouped data, and statistical analyses

For analyses and presentation of grouped data for various behaviours as a function of temperature, data obtained in individual experiments within the species were aligned against normalized range of the ramp (% T_LOE_, bottom axes in Figures 3 through 6, with 0 and 100% corresponding to initial temperature and T_LOE_, respectively, in a given experiment). Such treatment of the data was necessary to account for the differences in initial temperatures and in the LOE onset temperatures between experiments. Absolute ranges of temperature ramps in °C (top axes in Figures) are depicted as red plots above the traces, with red triangle symbols and horizontal error bars representing mean and SEM of the temperatures at the start of the ramp and at LOE averaged between respective numbers of experiments with each species.

Statistical significance of laterality bias was estimated by comparing lateralization indices at each 5% LOE temperature interval to a theoretical zero (random choice, 0% laterality) using one sample *t*-tests (Bisazza et al., 2000). Two-sample unequal variance two-tailed *t-*tests were used to determine statistical significance of changes in ventilation rate.

## RESULTS

### Locomotor responses to warming have alternating patterns

Fig. 1 depicts representative locomotor behaviours of *C. aceratus* and *N. coriiceps* elicited by acute warming, with marked transient bouts of increased locomotion of both species, interspersed with periods of reduced motility. Individual motility varies substantially within each species before and during warming, with Hb+ fish displaying more agility at all times. However, the alternating pattern of thermally induced changes in locomotion persists in all individuals with some species-specific trends in the onset, duration and velocity (Fig. S1).

**Figure 1.**
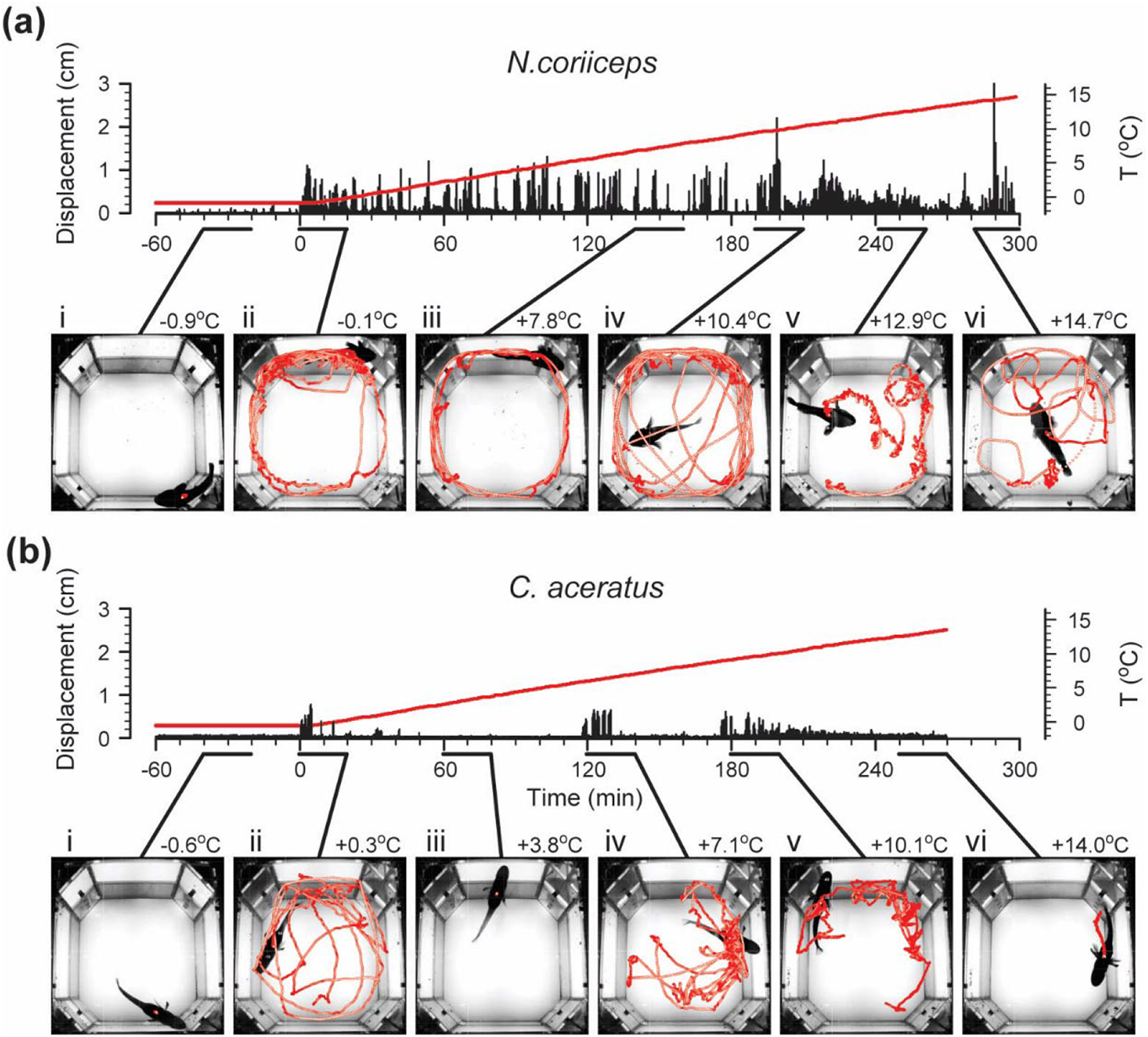
Representative locomotor responses of fishes to warming. (a) and (b) represent time plots of absolute instantaneous displacement (left axes) of a single specimen of *N. coriiceps* and of a single specimen of *C. aceratus*, respectively. Time “0” corresponds to the beginning of the temperature ramp; negative values correspond to the final sixty minutes of an overnight acclimation in the experimental tank. Red lines represent absolute temperature (right axes) of water, with the maximum corresponding to the LOE. Insets below time plots depict locomotion trajectories for 20 minute time periods at select temperatures (maximal temperatures during these episodes are shown next to the insets). (i) Baseline station-holding. (ii) Early responses to initial rise in temperature. (iii) First period of reduced motility. (iv) Intense locomotion and startle-like behaviours. (v) Second period of reduced motility with multiple startle-like behaviours. (vi) Onset of LOE.

The first locomotor responses to warming occur in all animals of both species with temperature elevations as little as 0.1°C, in agreement with early estimates of thermal sensitivity of fishes (Shelford and Powers, 1915; Bull, 1936). In striking contrast to sustained sedentary baseline behaviour (Figs. 1a (i) and 1b (i)), these manoeuvres manifest as yawing along the walls of the tank (Figs. 1a (ii) and 1b (ii)). Instantaneous velocities up to 0.8 body lengths per second (BL·s^−1^, see Materials and Methods for numerical BL data) in Hb+ and up to 0.4 BL·s^−1^ in Hb− fish are achieved predominately in the labriform swimming mode, with fore-aft rowing strokes of pectorals providing the thrust. Tail and trunk muscles are recruited only for occasional surfacings and short bouts of subcarangiform swimming, mostly evident in Hb+ fish.

Unable to perform thermoregulation by habitat selection in the tank, fishes respond to continued increase in temperature by reduction of motility (Figs. 1a (iii) and 1b (iii)). Although variable at the individual level, reduction of motility has marked commonalities within each species. Namely, as the temperature rises 1-1.5°C above initial, Hb− fish assume essentially motionless station-holding and maintain it for tens of minutes. In contrast to that, Hb+ fish generally continue to yaw along the tank walls, but gradually decrease the duration of swimming bouts. By +6°C, locomotion of Hb+ fish subsides, with station-holding episodes lasting up to ten minutes, interleaved occasionally with brief bouts of labriform swimming at velocities not exceeding 0.8 BL·s^−1^.

### Continued warming triggers startle-like responses

Conspicuous episodes of increased locomotion also occur between +5°C and +10°C in Hb− *C. aceratus*, and between +8.5°C and +12°C in Hb+ *N. coriiceps*, and manifest as intense swimming along the walls, accompanied by multiple surfacing events, particularly in Hb+ fish. Fishes use mainly the labriform swimming mode, achieving linear velocities up to ~1 BL·s^−1^ in Hb+ and ~0.6 BL·s^−1^ in Hb− animals (Fig. S1). In addition, characteristic only for *N. coriiceps*, they demonstrate several fast crossings of the tank, through its middle (Figs. 1a (iv) and 1b (iv)), using the subcarangiform propulsion with velocities reaching 2-3 BL·s^−1^ (Fig. S1a).

Notably, most of these manoeuvres involve distinctive patterns of body bending and subsequent turning. Namely, high-velocity swimming laps of Hb+ fish begin with a rotation of the head coincident with a typical contralateral tail bending, thus forming an S-shape (Figs. 2a and S3a). These highly dynamic S-bends are followed by very fast turns with angular velocities (V_a_) of ~1,000 degrees per second (deg·s^−1^). These manoeuvres, however, are scarce, with no more than five of them occurring in each animal. Turns of another type, recurrent in both species, begin with rotation of the head followed by ipsilateral bending of the tail, thus forming a C-shape (Figs. 2b, 2c, S3b and S3c). Such C-bend turns of *N. coriiceps* occur in a single stage, reaching maximal V_a_ of ~250 deg·s^−1^ (Fig. 2b). In contrast, multiple peaks of V_a_ below 100 deg·s^−1^ are evident during C-bend turns of *C. aceratus* (Fig. 2c). In both species, these manoeuvres are followed by either a short labriform swimming bout, or an unpowered glide of variable duration.

**Figure 2.**
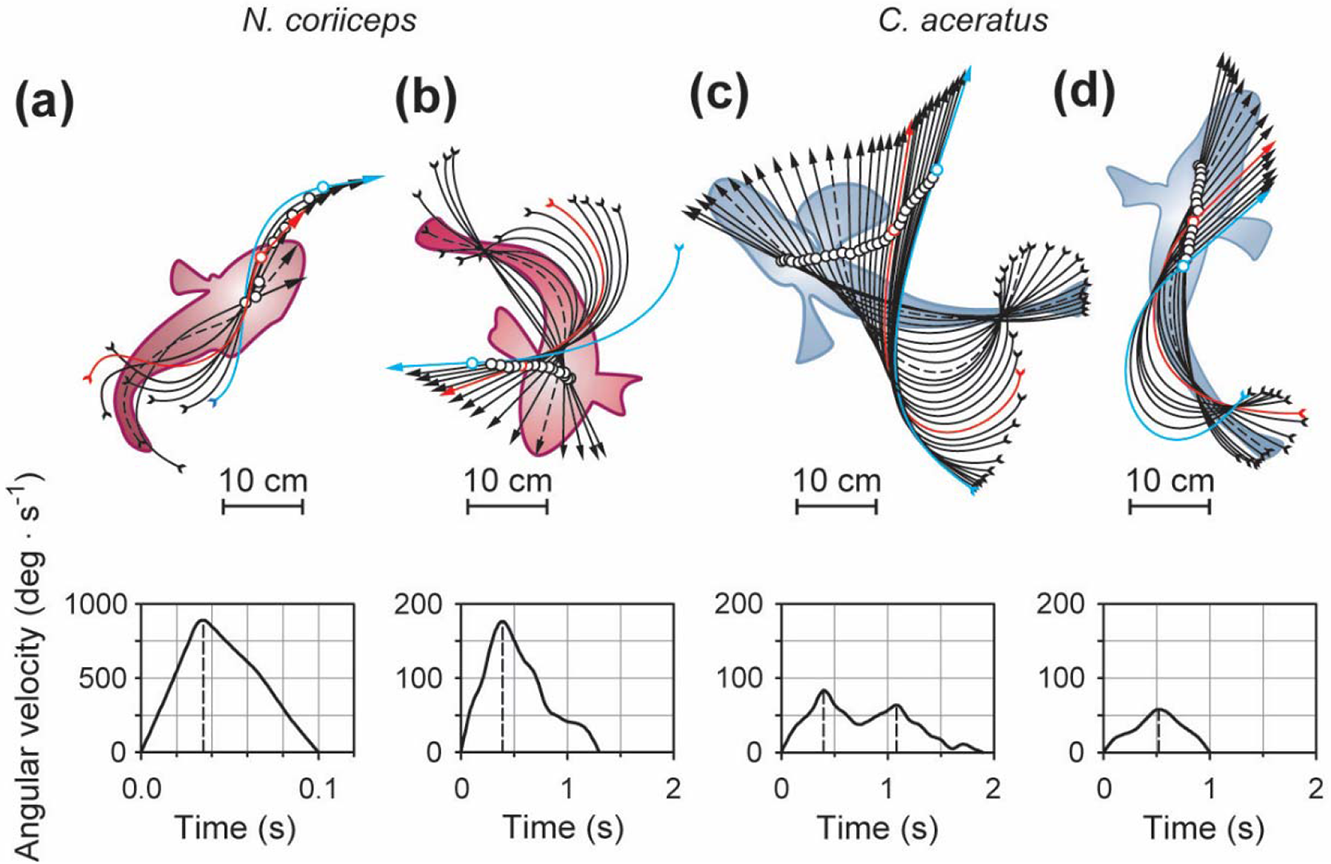
Repertoire of thermally induced startle-like manoeuvres. (a) Fast S-bend turn of *N. coriiceps* (33 ms intervals) at +11.1°C. (b) Intermediate velocity single-stage C-bend turn of *N. coriiceps* (100 ms intervals) at +12.4°C. (c) Slow multi-stage C-bend turn of *C aceratus* (100 ms intervals) at +8.5°C. (d) Withdrawal-like manoeuvre of *C aceratus* (100 ms intervals) at +11.5°C. Curved arrows and circles represent midlines and centres of mass of the fishes, respectively. Silhouettes with dashed midlines represent body-shapes and positions of fishes at maximal angular velocities during the turn stage of the manoeuvres (dashed vertical lines in the time plots of angular velocity). Red midlines indicate completion of the first stage of the manoeuvre (namely, the S- or C-bend proper) and transition to the second stage, swimming or gliding without change in direction (shown as blue midlines).

While these features are reminiscent of S- and C-starts in escape and startle behaviours reported in other fishes in response to external stimuli (Webb, 1976, Eaton et al., 1977; Domenici, 2010), 30 frames·s^−1^ sampling rate is suboptimal and may limit sufficient temporal resolution of body shapes. However, the midline and the silhouette of the fish in Fig. 2a have a discernible S-shape appearance at 33 ms, *i.e.*, at maximal angular velocity resolved during the turn stage. In addition, translation of the centre of the mass of *N. coriiceps* during manoeuvres depicted in Figs. 2a and 2b are consistent with early definitions of S-starts being characterised by displacement in line with the original body axis, and C-starts featuring large angles of turn (Webb, 1976; Webb, 1978; Eaton et al., 1977). Nonetheless, since the fish are not startled by an obvious stimulus, we term these thermally-induced manoeuvres “startle-like”, albeit differentiating initial S- and C-bends.

Unique to Hb− species, another type of thermally induced reactive behaviour features only slight rotation of the head with V_a_ below 50 deg·s^−1^, followed by retraction of the body (Figs. 2d and S3d). Observed in all five specimens of *C. aceratus*, it has the appearance of a slow backward movement of variable duration, and resembles “withdrawal” or “head retraction”, a startle response described in some other sedentary bottom-dwelling fishes with elongated bodies (Meyers et al, 1998; Ward and Azizi, 2004; Liu and Hale, 2014). While *C. aceratus* are indeed characterized by relatively high elongation ratios (see Materials and Methods for quantification and numerical data) and bottom-dwelling lifestyle, these thermally induced manoeuvres are much slower (~1 second in duration, Fig. 2d) than canonical withdrawals (which are over in 100-200 ms, Meyers et al, 1998; Ward and Azizi, 2004; Biermann et al 2004). Therefore, we term this behaviour “withdrawal-like”.

The repertoire and incidence of these startle-like behaviours of Hb+ and Hb− fishes demonstrate marked differences in thermal dependence (Fig. 3). While exact temperatures of the onset of S-bend manoeuvres of *N. coriiceps* vary between individual specimens, in each animal they occur within a relatively narrow interval of thermal change of ~1.5°C anywhere between +9.5 and +12°C. The intense swimming subsides after that in all Hb+ fish, followed by another period of reduced locomotion. However, they begin performing spontaneous C-bend manoeuvres, progressively increasing in incidence, and reaching up to 10 turns per minute between +12°C and +14°C (Figs. 3b and 1a(v)).

**Figure 3.**
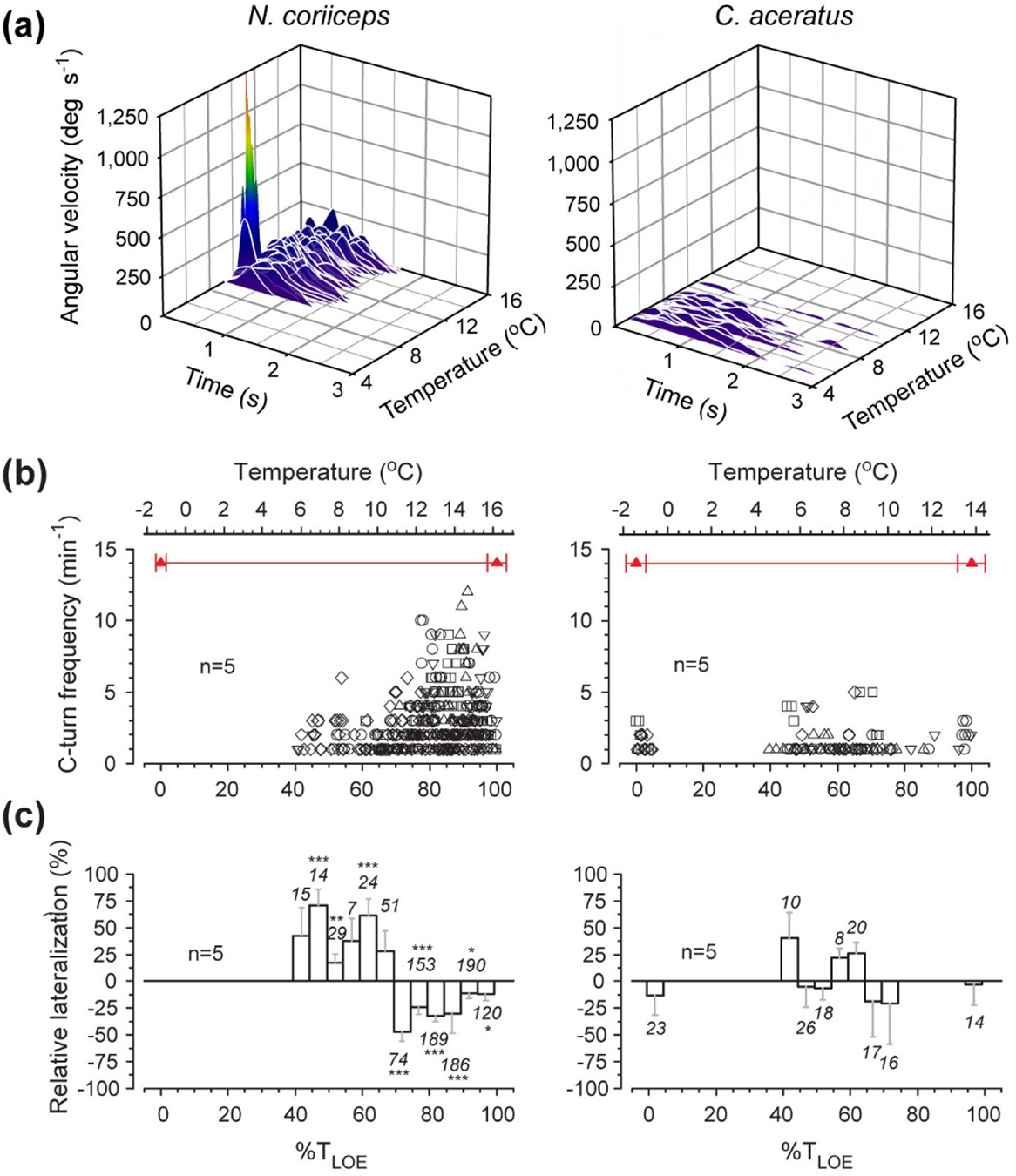
Startle-like behaviours as a function of temperature. a) Time plots of angular velocity (V_a_) during startle-like manoeuvres at respective temperatures in a single specimen of Hb+ and Hb− species. Rainbow colour code denotes the continuum of angular velocities from low (violet) to high (red) during fast S-bend turns (V_a_>500 deg·s^−1^, 9 events) and intermediate velocity single-stage C-bend turns in *N. coriiceps* (V_a_<250 deg·s^−1^, 137 events), and during slow multi-stage C-bend turns (V_a_<100 deg·s^−1^, 25 events) in *C. aceratus*. (b) Data points represent number of C-bend turns during each consecutive time interval one minute in duration in five specimens (depicted as different symbols) of each species. (c Relative lateralization (mean and SEM) of C-bend turns in 5% of T_LOE_ increments averaged between five specimens of each species. For conventions of quantification and presentation of grouped data as a function of temperature, see Materials and Methods. Numerals next to bars indicate numbers of events within each temperature increment in five animals. Asterisks denote statistical significance of laterality bias (*p*<0.05 - *; *p*<0.005 - **; *p*<0.001 - ***) at each temperature interval compared to a random choice (0% laterality) using one sample *t*-tests.

C-bend turns of *C. aceratus*, in contrast, are sporadic, and their frequency does not appear to change with temperature (Figs. 3b and 1b (v)). In this species, scarce withdrawal-like manoeuvres constitute substantial portion of motoric behaviours between +9°C and +12°C, occasionally interrupting extended episodes of station-holding. Nonetheless, startle-like behaviours persist in both fishes until the onset of LOE, when a short bout of erratic locomotion and surfacing occurs in all Hb+ and some Hb− animals (Figs. 1a (vi) and 1b (vi)).

Notably, below +11°C, startle-like C-bend turns of all *N. coriiceps* display a marked rightward preference (Fig. 3c, left graph). Above this temperature, this bias reverses, and the newly established leftward preference persists in subsequent C-bend turns of all five Hb+ fish.

With further warming, however, this lateralization gradually decreases and eventually disappears, when turns become scarce near the T_LOE_. Both the initial and reversed biases of turns of *N. coriiceps* are statistically significant (for *p*-values for each 5% LOE interval, see Fig. 3c, left graph), except when the behaviour just starts to manifest at 40% LOE, and at the point of right-left shift near 65% LOE. In contrast, none of five Hb− fish examined demonstrate any apparent bias of thermally induced C-bend manoeuvres (Fig. 3c, right graph).

### Coping with thermal stress involves ventilatory adjustments

In response to progressive aquatic hypoxia (Fig. S4) that is accompanied by escalating metabolic load associated with rising temperatures, fishes adjust their ventilation, as quantified in *post hoc* analyses of opercular movements in video recordings. Metrics of ventilation in three specimens of each species are shown in Fig. S5. At all temperatures, absolute ventilation rates of *N. coriiceps* are nearly two-fold higher than those of *C. aceratus* (Fig. S5a). However, because of high variability in locomotor activity (particularly between Hb+ specimens, Fig. S2) this difference in absolute ventilation rates between the Hb+ and Hb− species is not statistically significant (*p* > 0.8; two-sample two-tailed unequal variance *t-*test) except at their maxima (*p* < 0.02, two-sample two-tailed unequal variance *t*-test). Nonetheless, there are obvious similarities in the overall dynamics of changes in ventilation both within and between the species. Namely, after a small transient rise in ventilation frequency (*f_V_*) at the onset of warming ramp, once the temperature rises more than 2°C above initial, a steady increase in *f_V_* becomes evident in all animals (Figs. 4a and S5a). This hyperventilation persists even during episodes of relative quiescence, which suggests that it is not a consequence of locomotor effort. In both species, *f_V_* reach their maxima between +8 and +8.5°C, rising 2.5±0.9 times in *N. coriiceps* and 2.61±0.19 times in *C. aceratus* (normalized to the initial for each specimen; the increase is not statistically different between the species). In addition, near the maximum of *f_V_*, both fishes exhibit another well-known type of respiratory behaviour, aquatic surface respiration (Fig. S6). Ventilatory responses to further warming diverge between Hb+ and Hb− fishes, but there is substantial correlation of the overall changes in *f_V_* within the species, stronger in *C. aceratus* (*r* ranging from 0.88 to 0.96) than in *N. coriiceps* (*r* ranging from 0.40 to 0.71). Namely, after reaching the maximum, *C. aceratus* continue to hyperventilate until the 1–1.5°C prior to LOE, whereas *f_V_* of *N. coriiceps* declines precipitously within a rather narrow interval of temperature rise (~2°C) to a plateau at a lower, yet still relatively elevated, level.

**Figure 4.**
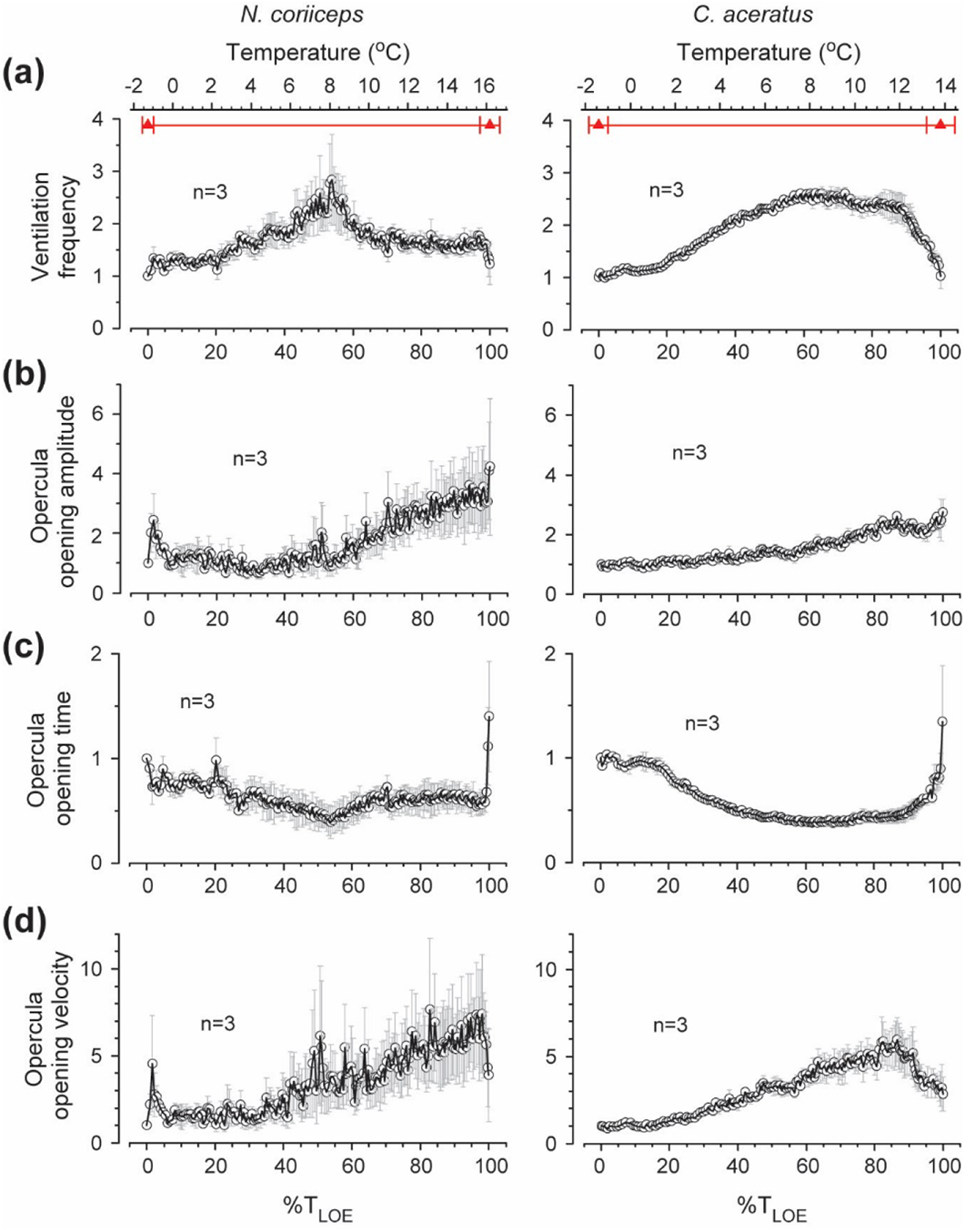
Thermally induced ventilatory responses. (a) Ventilation frequencies (*f_v_*). (b) Opercula opening amplitudes (OA). (c) Opercula opening times (time for opercula to transition from closed to open state, OT). (d) Opercula opening velocity (OV, calculated as OV = OA/OT). Data points and error bars in all plots represent means and SEM of each metric normalized to the initial value for each animal, and averaged for three specimens of each species. For conventions of quantification and presentation of grouped data as a function of temperature, see Materials and Methods.

Two other measures of opercular movements are the opening amplitude (OA) and the opening time (OT). With rising temperatures, OA increases progressively in both species, persistent till LOE (Fig. 4b). Hb− fish, however, recruit this adjustment at the onset of hyperventilation, whereas Hb+ fish employ it when *f_V_* approaches its peak. Thermally induced changes of OT, however, are biphasic, and nearly mirror the inverse of changes in *f_V_* in both species (Fig. 4c), reaching their minima nearly coincident with the maximal *f_V_* between +8 and +8.5°C. With further warming, *C. aceratus* maintain fast opercular openings (short OT) until the last 1–1.5°C prior to LOE, whereas in *N. coriiceps* they become slower (longer OT). Furthermore, during *f_V_*) plateaus, in both species, OT remains essentially unchanged, whereas OA steadily rises. This translates into continued growth of the ratio of these two measures of opercular movements, *i.e.*, opercula opening velocity (OV) (Fig. 4d), which keeps increasing while *f_V_* remains constant. When approaching LOE, at the temperatures above +15.5°C in *N. coriiceps*, and +11.5°C in *C. aceratus*, both *f_V_* and OV decline in both species followed by respiratory collapse.

### Continuous acute thermal stress triggers fanning and fin splay

Two other distinctive behaviours manifest in both species at the temperatures above +8°C. Both of them involve movements of pectoral fins with marked thermal dependences and species-specific differences in appearance.

One behaviour manifests as a cyclical fin movement in nearly stationary fishes, with no obvious relevance to locomotion, comparable to those reported in a variety of fishes during egg-guarding (Hancock, 1852) and termed “fanning”. In Hb− fish, it consists of anteroposterior undulations of large and flexible fan-shaped appendages, extended from the trunk (Fig. 5a, top panel, Supplementary Movies 1 and 2). These undulations have a mean frequency of ~1Hz (Fig. 5b), and occur in bouts lasting from tens to hundreds of seconds (Fig. 5c). Movements of less flexible pectorals of Hb+ fish present as co-mingled “sway” and “sweep” motions (Fig. 5a, bottom panel, Supplementary Movie 3). They are more than twice as slow as undulations in *C. aceratus* (Fig. 5b), and occur in short bouts (Fig. 5c).

**Figure 5.**
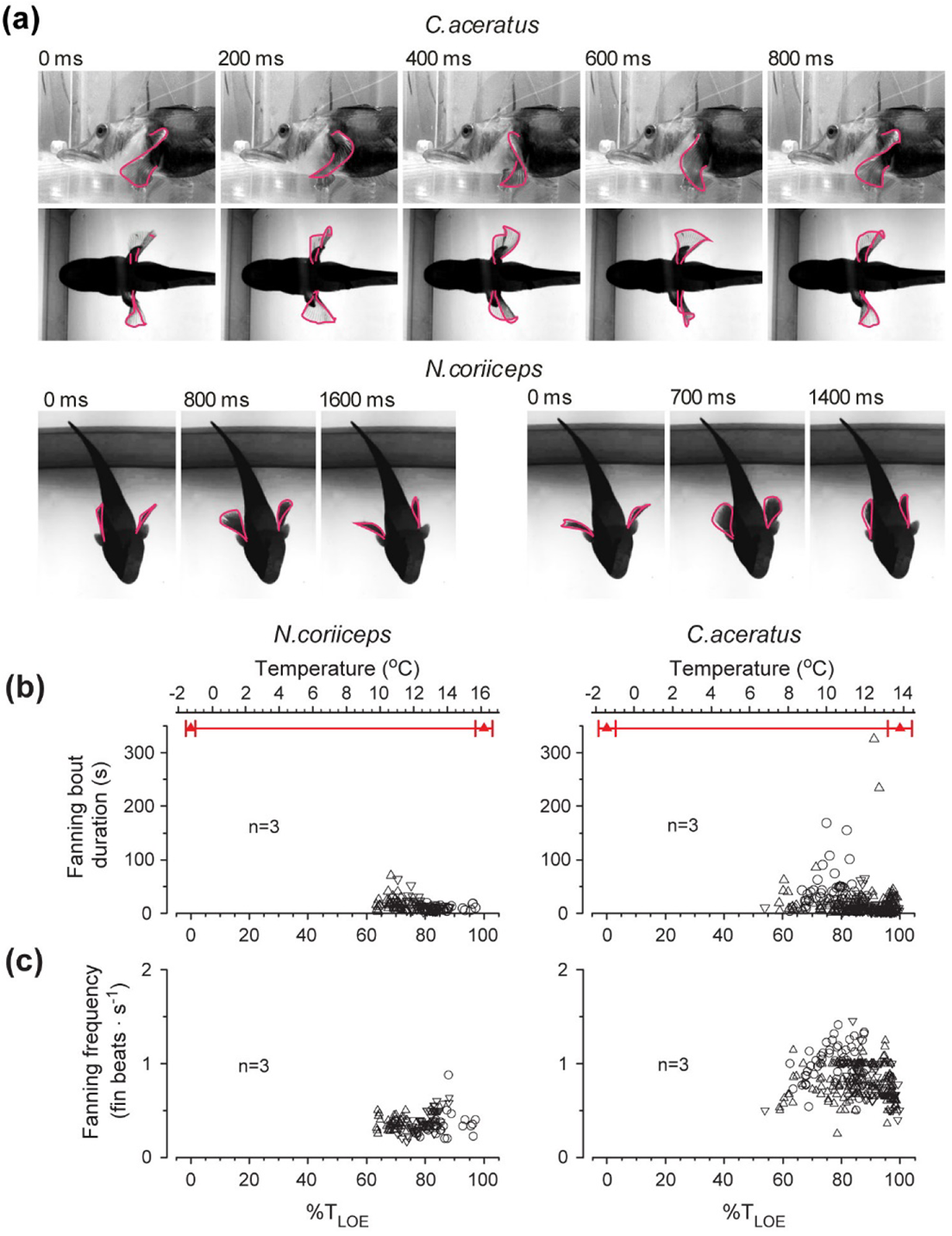
Acute warming elicits pectoral fin fanning. (a) Top panel depicts lateral (upper images) and ventral (lower images) view of one complete cycle (800 milliseconds in duration, every 6th frame shown) of undulatory fanning, typical for *C. aceratus*. Bottom panel depicts ventral view of one cycle of “sway” (fins adducted and abducted in a counterphase) fanning 1.6 s in duration (left set of images, every 24th frame is shown) and one cycle of “sweep” (fins adducted and abducted in a synphase) fanning 1.4 seconds in duration (right set of images, every 21st frame is shown), characteristic for *N. coriiceps*. (b) Temperature plots of duration of fanning bouts (episodes of uninterrupted continuous fanning at near constant frequency). (c) Temperature plots of fanning frequency (number of fin beats per second within a bout). For conventions of quantification and presentation of grouped data as a function of temperature, see Materials and Methods. Different symbols in (b) and (c) represent three specimens of each species.

In another behaviour, fishes spread their pectoral fins laterally, nearly perpendicular to the trunk, and maintain this position for a period of time (Fig. 6a). To the best of our knowledge, no comparable manoeuvre has ever been reported before, and we term it “splay” to depict the spreading of appendages.

**Figure 6.**
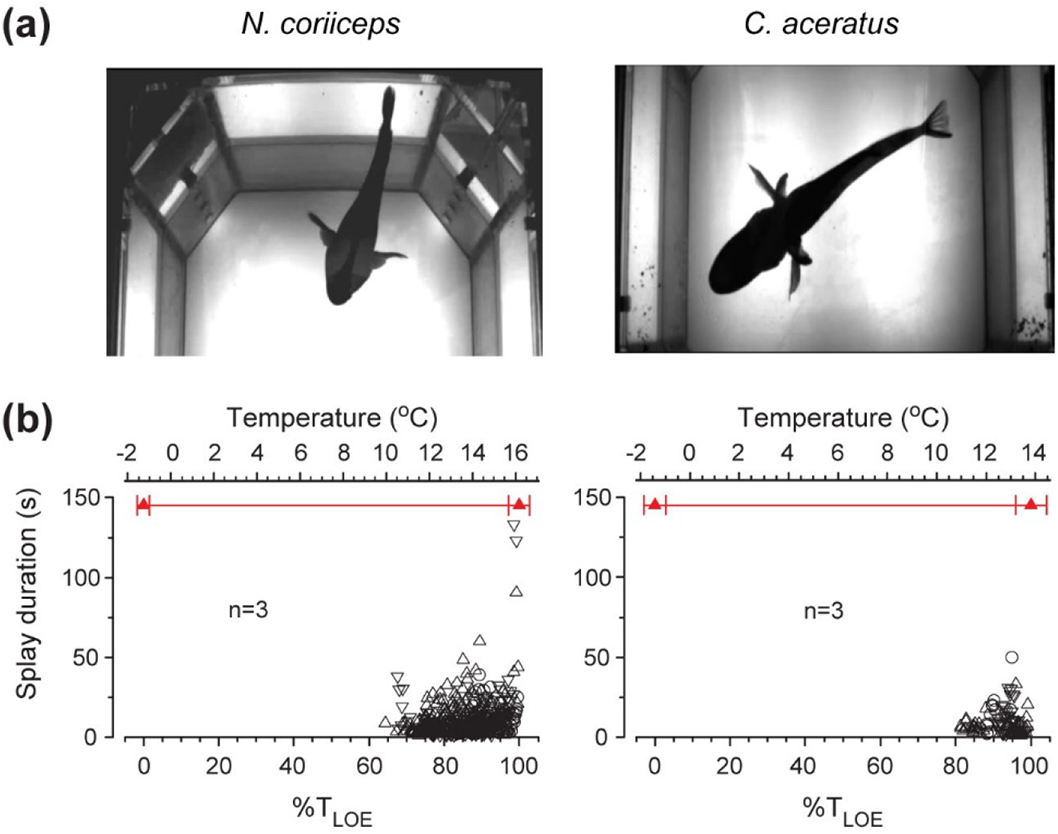
Fin splay behaviours at elevated temperatures. (a) Representative still images of splays in Hb+ and Hb− fishes. (b) Temperature plots of fin splay episode duration. Different symbols represent three specimens of each species. For conventions of quantification and presentation of grouped data as a function of temperature, see Materials and Methods.

Numerous splays of *N. coriiceps* are evident within relatively wide thermal range between +10°C and +16°C, increasing ~10 fold in occurrence by +13°C and up to 5-fold in duration by +16°C (Fig. 6b, left panel). Sporadic splays of *C. aceratus*, on the other hand, manifest between +9°C and +13°C, increasing ~4 fold in occurrence and up to 3-fold in duration (Fig. 6b, right panel).

### Patterned respiratory-locomotor coupling of N. coriiceps

Numerous fanning and splays of *N. coriiceps* are interspersed with C-bend turns, with certain stereotypy in the sequence of manoeuvres (Supplementary Movie 4), thus having features of “Fixed Action Pattern” (FAP) behaviour (Tinbergen, 1951). Namely, this behaviour manifests as copious repetitive Splay-Turn-Glide triplet sequences as often as five-six (up to ten near +14°C) times per minute (Fig 3B). The sequence of the three components remains invariant with increasing temperature, whereas the frequency of triplet occurrence, as well as duration of individual components appear to vary.

In addition, this patterned behaviour includes coordination between the movements of the opercula and rotation of the head. Namely, the acceleration and deceleration stages of head rotation during the C-bend turns are synchronized with the adduction and abduction phases of ventilatory opercular movements, respectively (Fig. 7).

**Figure 7.**
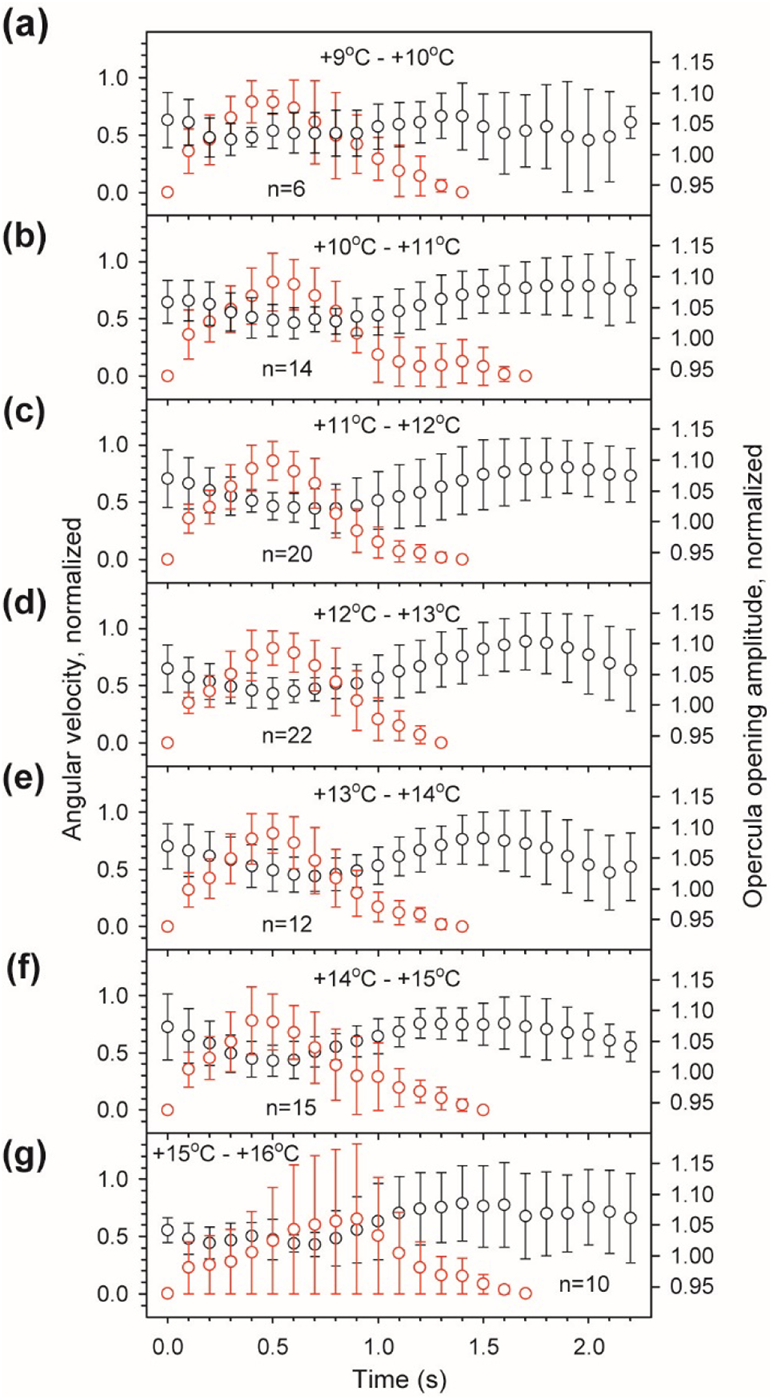
Respiratory-locomotor coupling in *N. coriiceps* during thermally induced C-bend manoeuvres. Data points and error bars represent mean and SD of angular velocity (red symbols) and opercula opening amplitude (black symbols), during C-bend turns of a single specimen of *N. coriiceps* (Cor #4 in Figs. S2 and S4 depicting locomotor and respiratory responses), normalized to their maximum and minimum values, respectively, and averaged for all successive turns within each 1°C interval of temperature change (see numbers in the graphs).

That is, C-bend turns routinely commence with opercula partially open, though undergoing adduction and closing near the peak of angular velocity. Opercula start opening again during angular deceleration and continue to open after the turn is complete, achieving the maximally open state when the fish is gliding. Such synchronization is evident in most C-bend turns at temperatures between +9°C and +15°C (Figs. 7a-7f), except within 1°C prior to LOE, when the synchrony is lost (Fig. 7g). Characteristically, these triplet manoeuvres synchronized with ventilatory movements of opercula involve only C-bend turns with V_a_ of ~250 deg·s^−1^, whereas V_a_ of S-bend turns (~1000 deg·s^−1^) is too fast to be coupled with opercula movements and gill ventilation.

## DISCUSSION

Major findings of our study are two-fold: 1) Antarctic Notothenoid fishes respond to acute ambient warming with an extensive repertoire of manoeuvres, most of which are observed in both Hb+ and Hb− species; 2) with some common tactics, behavioural strategies of Hb+ and Hb− fishes differ in terms of intensity, duration, thermal dependence and stereotypy of manoeuvres recruited. These commonalities and differences are discussed below in relation to possible physiological and ecological correlates.

Locomotion has been long recognized as the most obvious, and probably most universal, behavioural response of fishes to environmental stressors, particularly temperature (Shelford and Powers, 1915; Friedlander et al., 1976). From the rich repertoire of behaviours elicited in Hb− and Hb+ Antarctic fishes by acute warming, the earliest locomotor manoeuvres in response to the initial temperature rise may represent the attempts at ecologically relevant avoidance reactions. Also called habitat selection, this behavioural response has been considered the most essential thermoregulatory mechanism of ectotherms in a heterogeneous thermal environment (Crawshaw, 1979). However, essential for meaningful interpretation of the results of this study in the context of thermal tolerance and vulnerability to climate change, we must emphasize that our acute experiments in the laboratory setting differ from ecologically relevant venues in several aspects. First, although allowed a certain degree of freedom to express various behaviours, the fishes in the experimental tank are prevented from performing thermoregulation by habitat selection and are forced to thermoconform. Second, warming continues during this obligatory thermoconformation, further exacerbating the environmental stress. Third, the rate of temperature increase (~3°C per hour) is at least 10^5^ times faster than any current estimate of the rate of the Southern Ocean warming due to climate change (~3°C per 100 years, Vaughan et al., 2003; Cheng et al., 2019; Turner et al., 2016). Taking into account these aspects, we infer that most of the manoeuvres of the fishes observed in our experiments are aimed to physiologically mitigate detrimental effects of unavoidable acute warming.

We reason that reduced motility following initial presumed avoidance reaction may represent a strategy for conserving energy and preventing metabolic stress, similar to transitory “quiescent behaviour” observed in other fishes coping with progressive aquatic hypoxia (Nilsson et al., 1993; Schurmann and Steffensen, 1994). More prominent manifestation of this behaviour in Hb− fishes may suggest that lower oxygen-carrying capacity of their blood makes them more “prudent”.

With regard to startle behaviours, they are generally considered in a relatively narrow context of predator-prey interactions (Webb, 1976) or in relation to direct external stimuli (Eaton et al. 1977). Hence, environmental factors, such as temperature or DO_2_, are usually viewed in terms of their adverse effects on the success in avoiding predation (Domenici et al., 2007; Grigaltchik et al., 2012; Sánchez-García et al., 2019). Thermally induced startle-like manoeuvres in Antarctic fishes reported here for the first time suggest that similar behavioural patterns may be recruited not only in predator-prey interactions, but also in responses to environmental stress. Noteworthy in this regard, elevated temperature was reported to trigger escape-like turns in the African clawed frog tadpoles (Sillar and Robertson, 2009). Nonetheless, since the fish in our experiments are not startled by an obvious stimulus, and their movements during these responses are relatively slow, we reason that these thermally-induced manoeuvres are unlikely to represent *bona fide* startle behaviours (for recent definitions, see Domenici and Hale, 2019).

It is indeed possible that startle-like behaviours may result from the direct effects of elevated temperature on peripheral and/or central components of underlying neural circuits, activation of sensorimotor code and/or related motor pattern generating systems. Notably, all Notothenioids examined to date appear to lack (Eastman and Lannoo, 2004) the well identifiable T-shaped giant reticulospinal neurons in the hindbrain, the Mauthner cells, thought to initiate escape responses in most fishes (Zottoli and Faber, 2000; Eaton et al., 2001). Absence of obvious Mauthner cells, or presence of cells with deviant anatomy, has been reported in some fishes (Zottoli and Faber, 2000; Eaton et al., 2001), and their escapes were slower and significantly delayed (Greenwood et al., 2010). Our experiments do not provide any information about the latency of thermally induced startle-like manoeuvres since they appear to be spontaneous rather than evoked by an obvious external stimulus other than temperature. Notably, absolute values of maximal linear velocity (0.7 - 1.4 m·s^−1^) attained by *N. coriiceps* during thermally induced S-bend startle-like manoeuvres at +10 - +12°C are comparable with those during C-start escapes elicited by visual or tactile stimuli at 0°C in *N. neglecta* (1.28 m·s^−1^, Archer and Johnston, 1989) and in *N. coriiceps* (0.71 m·s^−1^, Franklin & Johnston, 1997). Thus, within a limited range of acutely elevated temperatures, Hb+ Antarcic fishes appear to be capable of maintaining high motor performance. No data on kinematics of startle behaviours of Hb− icefish is available for comparison.

The significance of lateralization of startle-like behaviours in *N. coriiceps* and its reversal is not immediately apparent. Preferred direction away from the startling stimulus has been seen in other fishes (Domenici, 2010). Lateralisation of barrier detours in consequent T-maze trials, on the other hand, appears to be poorly reproducible (Roche et al., 2020). Neither phenomenon, however, appears to have a relation to lateralization of spontaneous thermally induced turns of *N. coriiceps* observed in our experiments. Otherwise, aquatic hypoxia as well as elevated temperature have been demonstrated to alter both the direction and magnitude of behavioural laterality (Lucon-Xiccato et al., 2014; Allan et al., 2015), considered mainly in the context of predator-prey relationships. While exact mechanisms leading to these alterations are yet to be established, behavioural laterality is thought to reflect asymmetrical functional specialization of the brain across vertebrate species (Vallortigara & Rogers, 2005), with a rightward bias being controlled by the left hemisphere which is thought to govern routine behaviours, and a leftward bias controlled by the right hemisphere presumably responsible for emergency and stress reactions (Rogers, 2010).

Our findings of ventilatory adjustments while coping with progressive aquatic hypoxia and escalating metabolic load associated with rising temperatures are consistent with earlier observations in acutely warmed temperate (*e.g.*, Hughes and Roberts, 1970; Heath and Hughes, 1973) and some Hb+ Antarctic fishes (Fanta et al., 1989; Jayasundara et al., 2013). Furthermore, Jayasundara et al. (2013) report a bell-shaped thermal dependence of ventilation rate in another Hb+ Antarctic Notothenioid, *Trematomus bernacchii*, with a maximum near +8°C, comparable to our observations in *N. coriiceps*. Fast and substantial fall in *f_V_* observed after the ventilation maximum in Hb+ *N. coriiceps* (but not in Hb− *C. aceratus*) may result, at least in part, from autonomic splenic contraction which can rapidly boost the blood oxygen carrying capacity, similar to that seen following acute step-wise transfer of *Trematomus bernacchii* to +10°C (Davison et al., 1994). Noteworthy at this juncture, increased haematocrit has been reported in acutely warmed *N. coriiceps* at T_LOE_ (Beers and Sidell, 2011; Joyce et al, 2018b), although neither thermal dependence nor detailed physiology of the phenomenon are known. Otherwise, continued increase of ventilatory stroke volume (as deduced from measurements of OA, as a proxy metric) while maintaining nearly constant branchial pump suction (as deemed from measurements of OT, as an inverse proxy metric), evident in both Hb+ and Hb− species, suggests that during acute heat stress fishes employ additional, possibly more energetically advantageous, adjustments of ventilation by enhancing its efficacy (evident from changes in OV, as a proxy metric) without increasing *f_V_*. In addition, *N. coriiceps* may use synchronization of respiratory movements of opercula with head rotation during numerous C-bend turns to facilitate irrigation of the gills and thus increase respiratory efficiency via respiratory-locomotor coupling, comparable to that shown during undulatory swimming of a trout (Akanyeti et al, 2016).

As for fanning, earlier observations of this behaviour were made during egg-guarding, attributing it to parental care (Hancock, 1852), with ventilation of spawn being considered the main purpose under conditions of hypoxia and CO_2_ build-up in the nest (Van Iersel, 1953; Sevenster, 1961). Apparently widespread among Antarctic Notothenioids, egg-guarding does manifest uniparental fanning in some species (Daniels, 1979; Ferrando et al., 2014). In our experiments, however, fanning occurs in all specimens, male and female, in the absence of clutch, *i.e.*, under conditions that do not assume parental care. Based on the onset of fanning, nearly coincident with the maximum of *f_V_* and greatly intensifying during subsequent hyperventilation plateaus, we hypothesize that it constitutes an alternative respiratory behaviour in coping with thermal stress and progressive aquatic hypoxia. Supporting this hypothesis, continuous pectoral fanning was reported in captive *Trematomus loennbergii* in the McMurdo station aquarium (Eastman, 1993), also without clutch. Although the significance of this behaviour remained uncertain in this earlier report, it is plausible that some degree of hypoxia could exist in aquaria, even at low ambient temperature (*e.g.*, due to overcrowding). Specific roles of fanning in respiration of Hb− and Hb+ fishes, however, may differ. High frequency undulatory fanning in *C. aceratus* may facilitate cutaneous gas exchange, the role of which has long been discussed, particularly in the physiology of scaleless Hb− channichthyids (Hemmingsen, 1991; Eastman, 1993). Frequency of fanning in *N. coriiceps*, in contrast, is comparable with that of opercular beats, which may imply that in this species pectoral fin fanning may assist branchial pump.

Otherwise, all of these changes in respiratory behaviours of fishes observed in our experiments following the maximum of *f_V_* between +8 and +8.5°C (*i.e.*, after achieving the limit of ventilation effort) may be considered the manifestation of transition into “*pejus”* (from latin “worse”, Frederich and Pörtner, 2000) range. Albeit during this range they appear to mitigate, at least in part, the deleterious effects of progressive aquatic hypoxia and escalating metabolic load, respiratory collapse inevitably occurs in both species when all adjustments fail precipitously prior to LOE. It is not clear, however, if this collapse results from the limitations in cardiac function experienced by the fishes or is, conversely, their cause.

Regarding fin splays, the ethology of this newly described behaviour is not immediately apparent. We hypothesize that they may represent a possible contribution to cardiac output optimization during thermal stress, as they correspond to near maximal heart rates during thermal ramps (Joyce et al., 2018a). Specifically, the extension of appendages may move pectoral muscles and thus expand the pericardium which is, in fishes, attached to muscular elements of the pectoral girdle. In effect, the dimensions of pericardia of fishes are finite, thus imposing a limit on the maximal cardiac stroke volume. Indeed, a general observation is that fishes respond to warming by increasing their heart rate, whereas the stroke volume appears to be thermally insensitive (Eliason and Antilla, 2017). Some actively swimming fishes, including haemoglobinless *C. aceratus*, however, have demonstrated distinct increases in stroke volume in response to acute warming, particularly near the peaks of their heart rates (Gollock et al., 2006; Steinhausen et al., 2008, Mendonça and Gamperl, 2010; Joyce et al., 2018a).

Furthermore, surgical opening of the pericardium of a contracting *in situ* heart of haemoglobinless *Channichthys rhinoceratus*, resulted in a collapse of the ventricle (Feller et al., 1985), demonstrating the involvement of the intrapericardial pressure in the filling of the heart chambers in this species. On the other hand, in *N. coriiceps*, splays may work alongside respiratory-locomotor coupling during repetitive startle-like manoeuvres in Splay-Turn-Glide triplet FAP sequences, which persist in all Hb+ animals between ~12°C and ~15°C (over about one hour of warming ramp). On the other hand, while the sequence of the three components does not change with increasing temperature, the frequency of triplet occurrence and the duration of individual components within triplets vary, which is consistent with the concept of relative variability of FAPs (Schleidt, 1974). These newly described manoeuvres appear to involve multiple muscle groups (trunk, opercula, fins and, plausibly, heart) and thus may constitute more complex coordinated cardiac and respiratory mitigation of detrimental effects of acute heat stress.

Altogether, combinations of relative quiescence, changes in ventilatory effort and efficacy, respiratory-locomotor coupling during startle-like manoeuvres, fanning, and fin splay can all be reasoned as metabolic, respiratory, cardiac, and haematologic (latter only in Hb+ fish) accommodations, resulting in simultaneous concerted optimization of multiple vital functions. In the face of continuous warming, however, the capacity of all these physiological adjustments is limited, and they only provide for short-term compensation in extreme conditions. Furthermore, with an apparent multitude of physiological functions involved, neither the cause of the ensuant organismal failure, nor the apparent differences in tolerance of an acute sublethal thermal stress between the species can be attributed to a single organ or system.

Thus, our findings demonstrate considerable capacity of both Hb+ and Hb− Antarctic fishes for thermoconformation within a limited thermal range. These short-term behavioural and physiological adjustments may be imperative for transient migrations of eurybathic (Eastman, 2017) Notothenioids to favourable niches within a changing thermo- and oxycline for efficient habitat selection. Successful use of these new habitats under conditions of a lasting environmental change, however, should involve long-term behavioural and physiological adaptations, as well as ecological and evolutionary mechanisms (Huey et al, 2012; Pacifici et al., 2015). On the behavioural side, these adaptations may include adjustments of seasonal timing of life-history events (including reproduction) and biotic inter-species interactions (including predator-prey relationships). On the physiological side, development of long-term adaptations depends on the maintenance of successful, but usually rare, adaptive genetic variations (Brennan et al., 2019), which is contingent on the large census and effective population sizes (Pespeni et al., 2013). The latter may be particularly problematic for Notothenioids, many of which remain depleted after severe industrial over-harvesting (Kock, 1992) in the 1960-80s, with yet unclear prospects for population recovery (Belchier, 2013; Barrera-Oro et al., 2017). That is, without rational science-based management of fisheries in the Southern Ocean (Brooks et al., 2018), unregulated anthropogenic interventions have the potential to produce irreversible damage to this ecosystem, irrespective of how well species adapt to the detrimental effects of climate change.

## DATA AVAILABILITY

The published article includes all datasets generated or analysed during this study. Original videos used for tracking analyses are available from the corresponding authors on request.

## ABBREVIATIONS

BL·s^−1^: body lengths per second DO_2_ - dissolved oxygen
FAP: Fixed Action Pattern *f_V_* - ventilation frequency Hb− - haemoglobinless
Hb+: haemoglobin expressing
LOE: loss of equilibrium
OA: opercula opening amplitude
OT: opercula opening time V_a_ - angular velocities

## ACKNOWLEDGEMENTS

We are indebted for superb logistic support to the masters and crew of the Antarctic Research and Supply Vessel *Laurence M. Gould*, and to the winter support personnel of the United States Antarctic Program Palmer Station. Animal acquisition and maintenance during field work for this study represents a collaborative effort of ourselves and Dr. Elizabeth L. Crockett (Ohio University, USA), Dr. Kristin M. O’Brien (University of Alaska Fairbanks, USA), Dr. Stuart Egginton (University of Leeds, UK), Dr. Anthony P. Farrell (University of British Columbia, Canada), Dr. Theresa J. Grove (Valdosta State University, USA), Ms. Amanda M. Biederman (Ohio University, USA) and Ms. Elizabeth Evans (Ohio University, USA). Funding for the field work was provided by the National Science Foundation grant ANT 1341602 to Dr. Elizabeth L. Crockett (Ohio University, USA). We are also indebted to Dr. Steve Zottoli for his critical suggestions on the manuscript. We also thank Dr. Chris Darby, the United Kingdom Chief Scientist to the Commission for the Conservation of Antarctic Marine Living Resources (CCAMLR), and Dr. Mark Belchier, the originator of the data, for permission to access and use the CCAMLR document WG-FSA 13/26. Funding for the field work was provided by the National Science Foundation grant ANT 1341602 to Dr. Elizabeth L. Crockett (Ohio University, USA). Acquisition of equipment for behavioural experiments and all other activities throughout the study including the time and effort of Drs. Ismailov, Scharping and Friedlander were supported by the Fralin Biomedical Research Institute at VTC operational funds.

## DECLARATION OF INTEREST

None.

## AUTHOR CONTRIBUTIONS

Conceptualization - M.J.F.; Overall Methodology - M.J.F., I.I.I., J.B.S.; Equipment Custom Design - I.I.I., I.E.A., J.B.S.; Experiments at Palmer Station - I.I.I., J.B.S; Data analyses - I.I.I., I.E.A., J.B.S; Writing of the MS: original draft - I.I.I., I.E.A.; review and editing - I.I.I., I.E.A., J.B.S; M.J.F.

## SUPPLEMENTARY FIGURES

**Figure S1.**
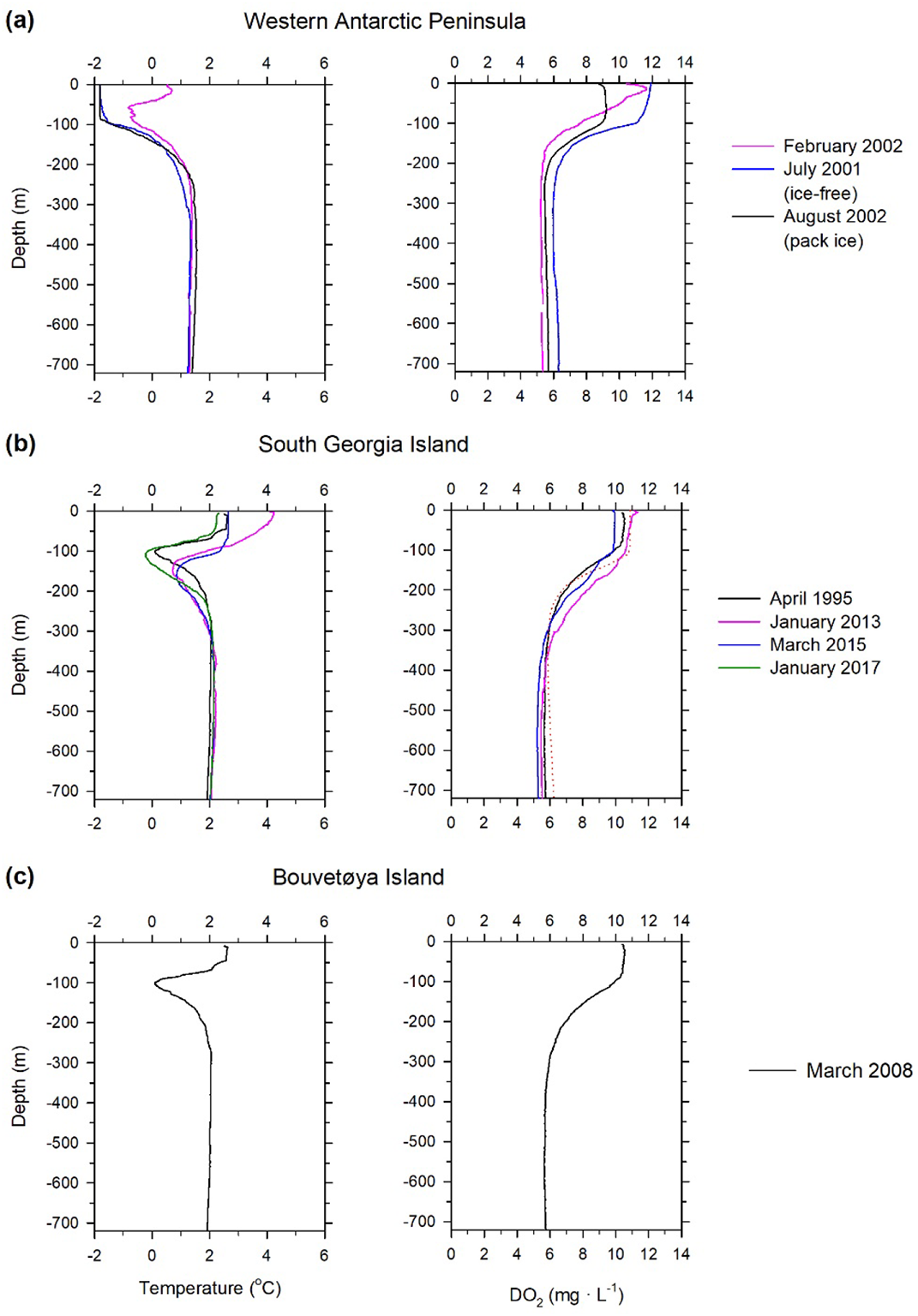
Metadata analysis of vertical temperature and DO_2_ profiles in areas of historic natural habitat of *N. coriiceps* and *C. aceratus*. (a) – Vicinity of the Western Antarctic Peninsula; Data sources: February, 2002 - Fritsen, C. (2011) CTD data from the Cruise CD-ROMS on ARSV Laurence M. Gould (LMG0201A, LMG0203, LMG0205, LMG0302, LMG0104) in the Southern Ocean from 2001-2003 (SOGLOBEC project). Biological and Chemical Oceanography Data Management Office (BCO-DMO). Dataset version 2011-04-07. Subset LMG0201 (Cast ## A1, A2, B2). http://lod.bco-dmo.org/id/dataset/2361; accessed 3/14/2017; July, 2001 and August, 2002 - Klinck, J.M. and Hofmann, E.E. (2003) Processed standard depth (pressure) CTD data from SOGLOBEC survey cruises on RVIB Nathaniel B. Palmer - NBP0103; NBP0104; NBP0202; NBP0204 - in the Southern Ocean from 2001-2002 (SOGLOBEC project). Biological and Chemical Oceanography Data Management Office. Dataset version 2003-06-13. Subsets NBP0104 and NBP0204. http://lod.bco-dmo.org/id/dataset/2360; accessed 3/17/2017. Data on pack ice situation in August 2002 - U.S. Southern Ocean GLOBEC Report No. 8. Report of RVIB Nathaniel B. Palmer Cruise NBP02-04 to the Western Antarctic Peninsula, 31 July to 18 September 2002. http://www.ccpo.odu.edu/Research/globec/main_cruises02/nbp0204/menu.html (b) – Vicinity of the South Georgia Island; Data source - British Oceanographic Data Centre, Natural Environment Research Council, UK. April, 1995, January, 2013 and March, 2015 and January, 2017 curves represent averages of STD/CTD data collected during the RRS James Clark Ross cruise JR19950320 (JR10) within The UK World Ocean Circulation Experiment (WOCE) Project (BODC ID 1011019, 1011020, 1011032, 1011044 and1011068); during RRS James Clark Ross cruise JR20130109 (JR274) within UK Ocean Acidification Research programme (BODC ID 1147675, 1147687, 1147699 and 1147706); during RRS James Clark Ross cruise JR20150309 (JR272D, JR310) within British Antarctic Survey Long Term Monitoring and Survey programme (BODC ID 1814444, 1814456, 1814468, 1814481 and 1814493); and during RRS James Clark Ross cruise JR16004 within the Ocean Regulation of Climate by Heat and Carbon Sequestration and Transports (ORCHESTRA) project of the Natural Environment Research Council, UK (BODC ID 1836882, 1836894, 1836901 and 1836925), respectively. January 2017 dataset does not have usable DO_2_ data. (c) – Vicinity of the Bouvetøya Island; Data source - British Oceanographic Data Centre, Natural Environment Research Council, UK. Curves represent averages of STD/CTD data collected during the Marion Dufresne cruise MD166 (BONUS-GOODHOPE, GIPY04) within the GEOTRACES project of the University of Western Brittany, France (BODC ID 1105470, 1105494, 1105433). Temperature of surface waters varies between locations during austral summer. Surface water DO_2_ in the vicinity of the Western Antarctic Peninsula does not vary much between seasons, but depends on pack ice cover (compare July, 2001 and August, 2002 DO_2_ profiles in **a,** right panel). No STD/CTD data are available for austral winter in the vicinity of South Georgia and Bouvetøya islands. Preferred bathymetric ranges of *Nototheniidae* and *Channichthyidae* (<200 metres for *N. coriiceps* and >100-200 metres for *C. aceratus*; Hureau, 1985), correspond to pronounced thermo- and oxycline in all locations.

**Figure S2.**
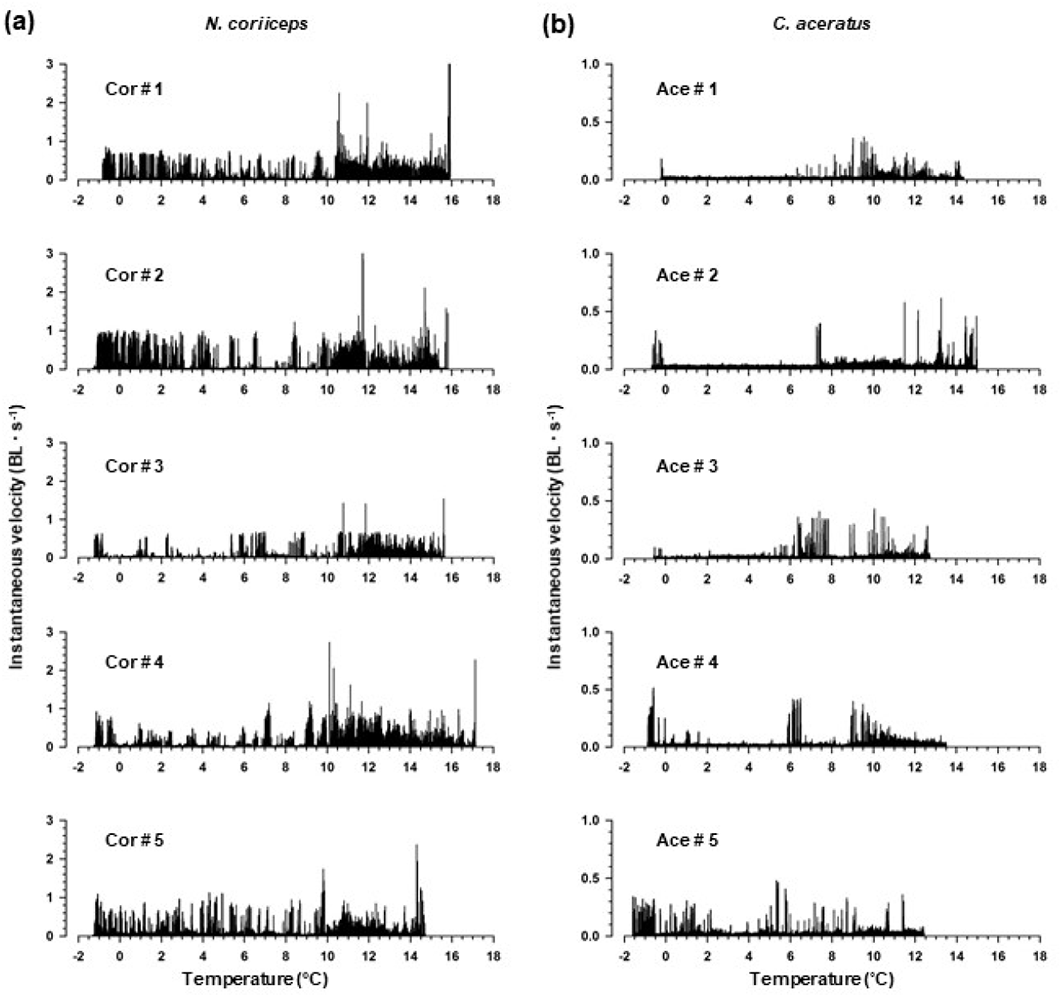
Locomotor responses to warming in different specimens of *N. coriiceps* and *C. aceratus*. Traces are temperature plots of instantaneous (30 Hz sampling rate) velocity in individual experiments with (a) five specimens of N. coriiceps and (b) five specimens of C. aceratus calculated from absolute displacement of the centre of the mass of the fish in each consecutive sampling interval (video frame), normalized for body length (BL) of the specimen. Overall, N. coriiceps are more motile than C. aceratus (note a difference in scales of ordinates). Note that initial temperatures vary between experiments due to changing weather conditions in the Arthur Harbour. However, all animals of both species exhibit presumed avoidance behaviours within initial rise in temperature of as little as 0.1°C, followed by quiescence and reduced motility (sluggishness in N. coriiceps and motionless in C. aceratus). Onset and amount of intense locomotion and startle-like behaviours vary between individual animals within the species. During these manoeuvres, N. coriiceps achieve instantaneous velocities up to 2.5 BL·s-1. When approaching loss of equilibrium, all animals of both species reduce motility, but perform multiple startle-like manoeuvres. Prior to the onset of LOE, all N. coriiceps and some specimens of C. aceratus exhibit fast erratic swimming.

**Figure S3.**
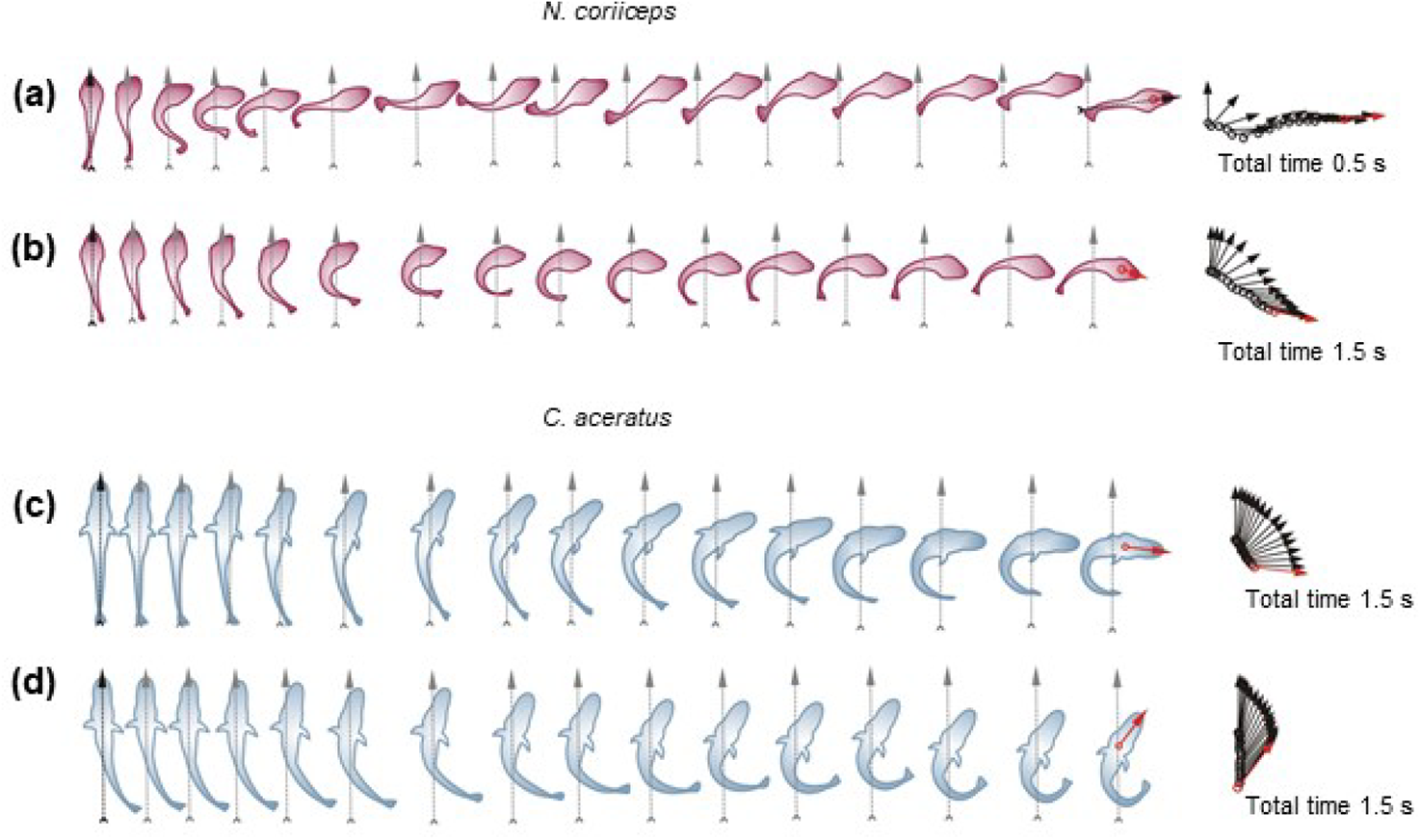
Axial movements during temperature induced startle-like responses in *N. coriiceps* and *C. aceratus* in ventral view (pectoral fins not shown) (a) *N. coriiceps* during an S-bend manoeuvre shown at 33 ms intervals (at +11.05°C) (b) *N. coriiceps* during a C-bend turn shown at 100 ms intervals (at +12.4°C) (c) *C. aceratus* during a C-bend turn shown at 100 ms intervals (at +8.5°C) (d) *C. aceratus* during a withdrawal (head retrieval) shown at 100 ms intervals (at +11.5°C) The grey vertical arrows indicate the position of the midline at the beginning of turning manoeuvres. Arrow diagrams on the right depict of the head orientation during turns, at time intervals indicated above. Each arrow connects the position of the fish nose with the centre of mass. Red arrows indicate final direction of locomotion. Note that C-bend turns of *N. coriiceps* are almost twice faster than those of *C. aceratus (cf.* (b) and (c)).

**Figure S4.**
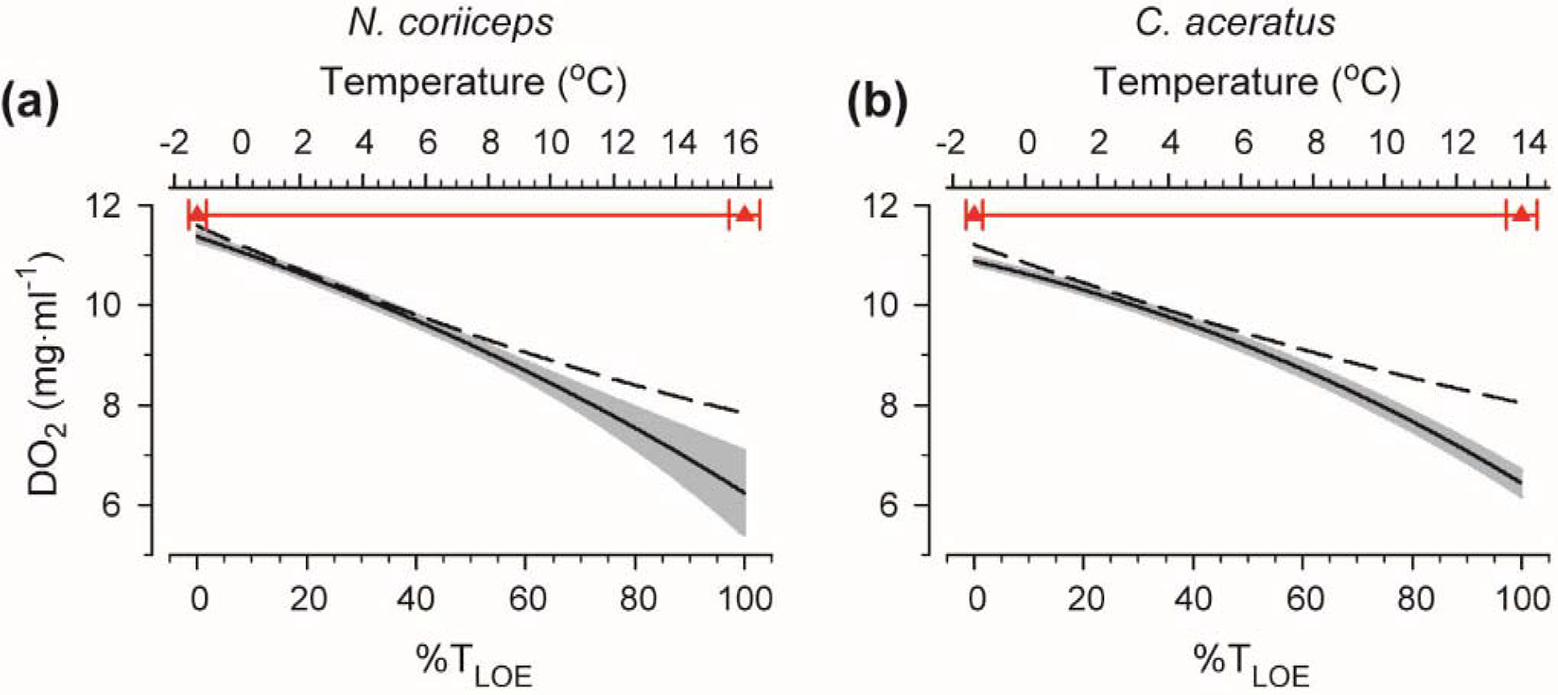
Progressive aquatic hypoxia concomitant with warming of water in the experimental tank. Solid black lines and grey error bars in the graphs represent means and SEM of absolute DO_2_ measured simultaneously with temperature ramps, averaged between five experiments of each species. Dashed lines represent theoretical temperature plots of DO_2_ calculated using Henry’s law, corrected for salinity. For conventions of analyses and presentation of grouped data as a function of temperature, see legend for Fig. 2 and Materials and Methods. Briefly, to account for the differences in initial temperatures and in LOE onset temperatures between experiments, for analyses and presentation of grouped data for various behaviours as a function of temperature, data obtained in individual experiments within the species were aligned against each other and against normalized range of the ramp (% T_LOE_, bottom axis), with 0 and 100% representing initial ambient temperature and temperature at LOE in each respective experiment. Absolute ranges of temperature ramps in °C (top axis) are depicted as red plots, with red triangle symbols and horizontal error bars representing mean and SEM of temperatures at the start of the ramp and at LOE averaged between five experiments with each species. Whether deviation of empirical DO_2_ temperature plots from theoretical represents metabolic O_2_ consumption by the fishes, or from nonlinearities of our measurements is open to conjecture since 1) manufacturer of the instrument does not imply calibration of the probe at temperatures below 0°C; 2) in the absence of conductivity measurements made during our experiments, we used theoretical correction for salinity.

**Figure S5.**
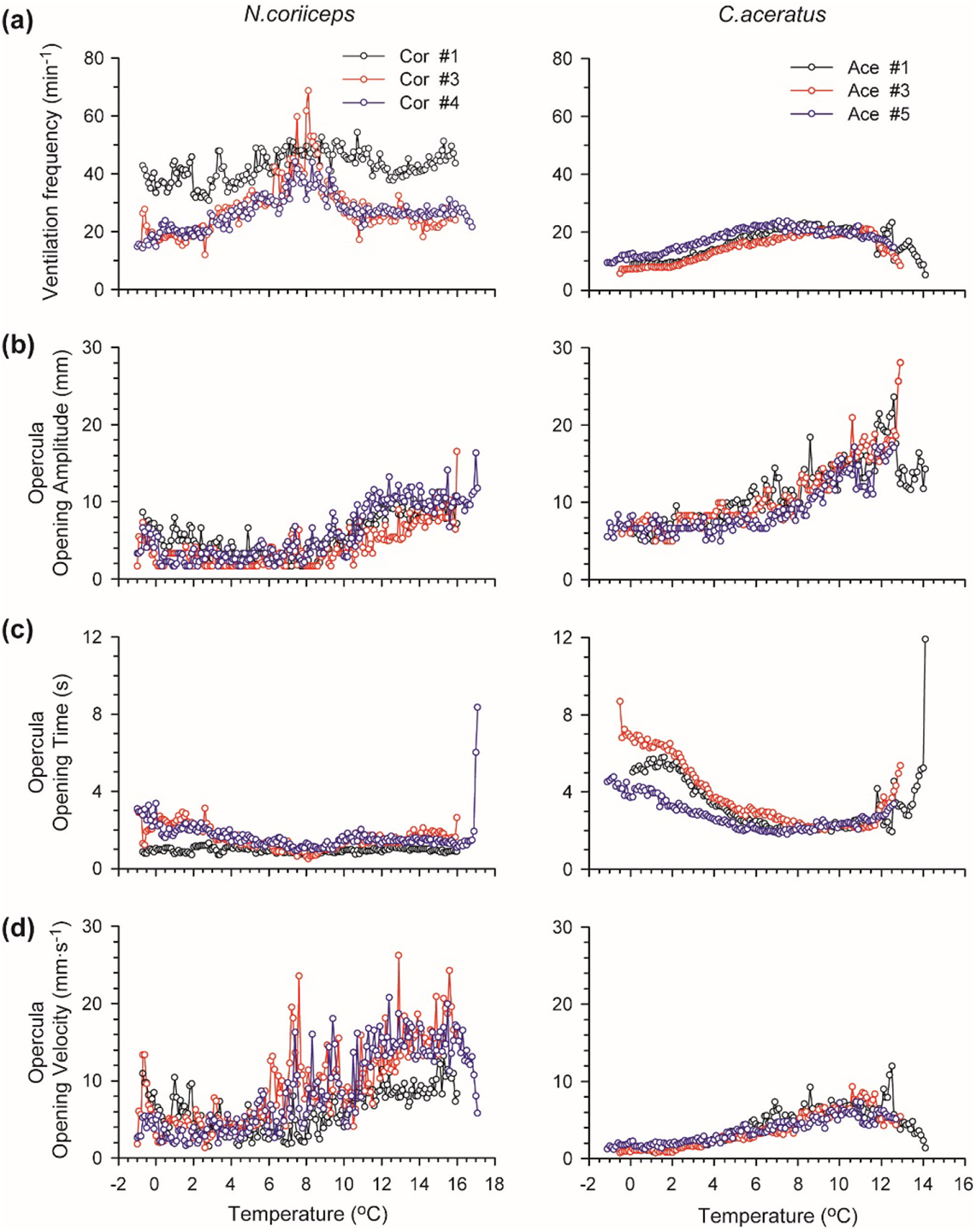
Ventilatory responses to warming in different specimens of *N. coriiceps* and *C. aceratus*. In response to warming, all fishes transiently increase ventilation rates (fV), amplitudes of opercula opening (OA), but decrease opening time of opercula (OT). Data points represent respective metrics in three specimens of each species (## in legends correspond to individual experiments shown in Fig. S1). Note high fV in Cor # 1 (black trace in top left panel) throughout entire experiment, likely due to overall higher locomotor activity, possibly masking the effects of temperature.

**Figure S6.**
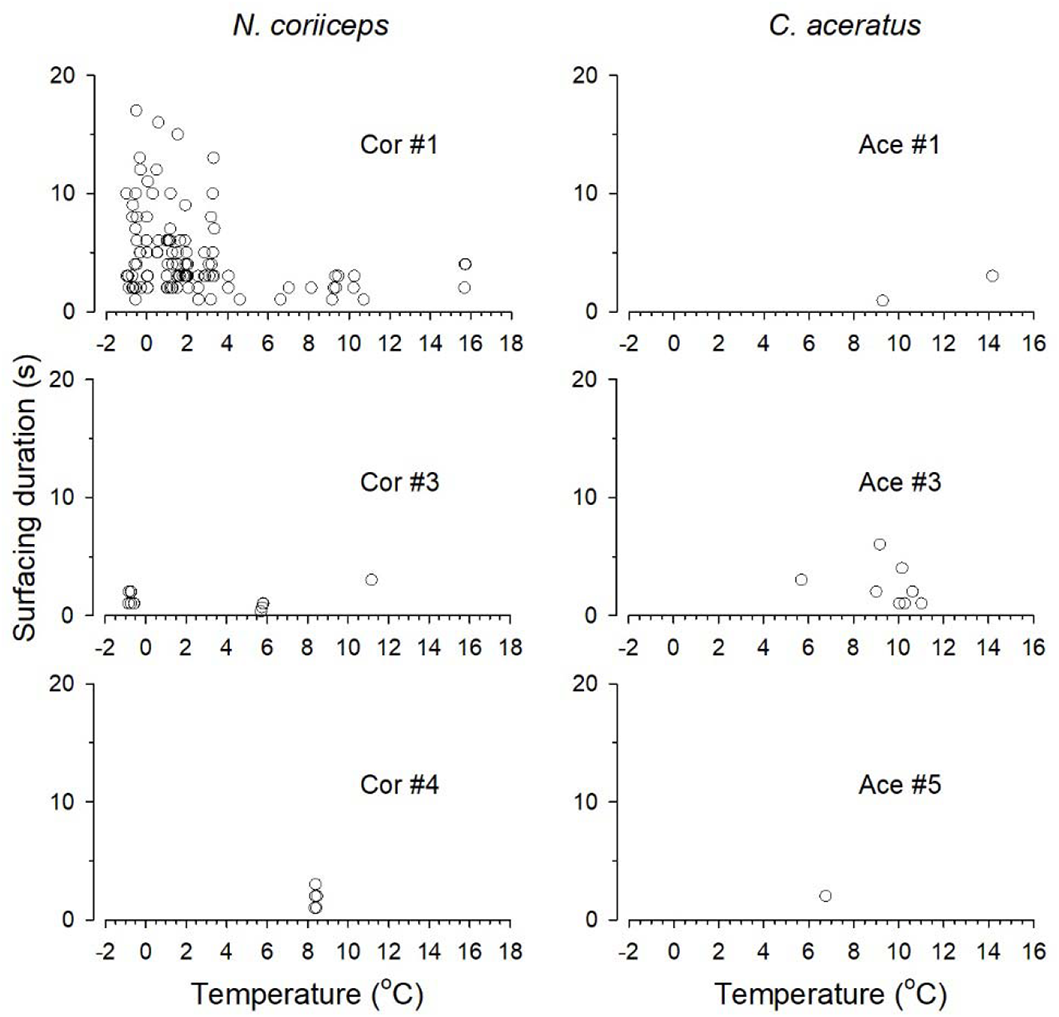
Aquatic surface respiration of *N. coriiceps* and *C. aceratus* as a function of temperature. Data points represent time spent by three specimens of each species at water-air interface in a form of continuous uninterrupted surfacing bouts, labelled corresponding to legends for Figs. S1 and S4. Occurrence of surfacing bouts in N. coriiceps is highly variable between the specimens, ranging from >10 to <100 in number, from <0°C to >+6°C in the onset temperatures, and from 1 to 17 seconds in duration. Occurrence of surfacing in C. aceratus is less variable, ranging from one to 8 in number, with all but two events occurring at the temperatures between +8°C and +12°C, and all lasting from 1 to 7 seconds in duration. Surfacing of C. aceratus and short (< 7 seconds in duration) surfacing of N. coriiceps seen at the temperatures between +6°C and +12°C may represent aquatic surface respiration behaviour, triggered by progressive aquatic hypoxia, inherent to diminished solubility of gases in water at elevated temperatures. Ethological understanding of numerous relatively long (lasting up to 17 seconds) surfacing elicited in some (but not all) Hb+ fishes at low to moderate levels of hypoxia is not immediately apparent. We hypothesize that they may reflect the stress elicited by warming in fish deprived from a choice of habitat/temperature in the tank.

## LIST OF SUPPLEMENTARY MOVIES

**Supplementary Movie 01** - *C. aceratus* fanning at +10.5°C – ventral view https://drive.google.com/file/d/1EJxThzRoSFLgZIesDaAdu47k4BL6uPLK/view?usp=sharing

**Supplementary Movie 02** - *C. aceratus* fanning at +10.5°C – side view https://drive.google.com/file/d/1-PoIQMMfG_qf0srpxVl7IMepgVQG1e8X/view?usp=sharing

**Supplementary Movie 03** - *N. coriiceps* fanning at +11.2°C https://drive.google.com/file/d/1R_bQb19adHFC4cQpRWLNn2ON 50cBhY/view?usp=sharing

**Supplementary Movie 04** - *N. coriiceps* splays and FAPs at +14.7°C https://drive.google.com/file/d/1J4U-qU2f2QJnIHH4vnC_6fLkILjWBJDI/view?usp=sharing

## Notes

### Competing Interest Statement

The authors have declared no competing interest.

### Summary of Updates

Revised format of the Reference List.

## REFERENCES

1. Akanyeti, O., Thornycroft, P. J., Lauder, G.V., Yanagitsuru, Y.R., Peterson, A.N. and Liao, J.C. (2016). Fish optimize sensing and respiration during undulatory swimming. Nat. Comm. 7, 11044 https://doi.org/10.1038/ncomms11044.

2. Allan, B.J., Domenici, P., Munday, P.L. and McCormick, M.I. (2015). Feeling the heat: the effect of acute temperature changes on predator-prey interactions in coral reef fish. Cons. Physiol. 3, cov011 https://doi.org/10.1093/conphys/cov011.

3. Archer, S.D. and Johnston, I.A. (1989). Kinematics of labriform and subcarangiform swimming in the Antarctic fish *Notothenia neglecta*. J. Exp. Biol. 143, 195–210.

4. Barker, P.F. and Thomas, E. (2004). Origin, signature and palaeoclimatic influence of the Antarctic Circumpolar Current. Earth Sci. Rev. 66, 143–162 https://doi.org/10.1016/j.earscirev.2003.10.003.

5. Barrera-Oro, E., Marschoff, E. and Ainley, D. (2017). Changing status of three notothenioid fish at the South Shetland Islands (1983–2016) after impacts of the 1970–80s commercial fishery. Polar Biol. 40, 2047–2054 https://doi.org/10.1007/s00300-017-2125-0.

6. Beers, J.M. and Sidell, B.D. (2011). Thermal tolerance of Antarctic notothenioid fishes correlates with level of circulating hemoglobin. Physiol. Biochem. Zool. 84, 353–362 https://doi.org/10.1086/660191.

7. Belchier, M. (2013). *Decadal trends in the South Georgia demersal fish assemblage*. SCCAMLR-WG-FSA-13/26, CCAMLR, Australia, Hobart, 26 pp., https://www.ccamlr.org/en/wg-fsa-13/26.

8. Bierman, H.S., Schriefer, J.E., Zottoli, S.J. and Hale, M.E. (2004). The effects of head and tail stimulation on the withdrawal startle response of the rope fish (*Erpetoichthys calabaricus*). J. Exp. Biol. 207, 3985–3997 https://doi.org/10.1242/jeb.01228.

9. Bisazza, A., Facchin, L., Pignatti, R. and Vallortigara, G. (1998). Lateralization of detour behavior in poeciliid fish: the effect of species, gender and sexual motivation. Behav. Brain Res. 91, 157–164 https://doi.org/10.1016/S0166-4328(97)00114-9.

10. Bisazza, A., Cantalupo, C., Capocchiano, M. and Vallortigara, G. (2000) Population lateralisation and social behaviour: a study with 16 species of fish. Laterality 5, 269–284 https://doi.org/10.1080/713754381.

11. Brennan, R.S., Garrett, A.D., Huber, K.E., Hargarten, H. and Pespeni M.H. (2019). Rare genetic variation and balanced polymorphisms are important for survival in global change conditions. Proc. R. Soc. B 286, 20190943 https://doi.org/10.1098/rspb.2019.0943.

12. Brooks, C.M., Ainley, D.G., Abrams, P.A., Dayton, P.K., Hofman, R.J., Jacquet, J. and Siniff, D.B. (2018). Antarctic fisheries: factor climate change into their management. Nature 558, 177–180 https://doi.org/10.1038/d41586-018-05372-x.

13. Bull, H.O. (1936). Studies on conditioned responses in fishes. Part VII. Temperature perception in teleosts. J. Mar. Biol. Ass. 21, 1–27 https://doi.org/10.1017/S0025315400011176.

14. Cheng, L., Abraham, J., Hausfather, Z. and Trenberth, K.E. (2019). How fast are the oceans warming? Science 363, 128–129 https://doi.org/10.1126/science.aav7619.

15. Clarke, A., Barnes, D.K.A., Bracegirdle, T.J., Ducklow, H.W., King, J.C., Meredith, M.P., Murphy, E.J. and Peck, L.S. (2012). The impact of regional climate change on the marine ecosystem of the Western Antarctic Peninsula. In Antarctic ecosystems: an extreme environment in a changing world (ed. A.D. Rogers, N.M. Johnston, E.J. Murphy, and A. Clarke), pp. 91−120. Oxford, UK, Wiley-Blackwell https://doi.org/10.1002/9781444347241.ch4.

16. Cocca, E., Ratnayake-Lecamwasam, M., Parker, S.K., Camardella, L., Ciaramella, M., di Prisco, G. & Detrich, H.W., 3rd (1995). Genomic remnants of alpha-globin genes in the hemoglobinless Antarctic icefishes. Proc. Natl. Acad. Sci. USA 92, 1817–1821 https://doi.org/10.1073/pnas.92.6.1817.

17. Constable, A.J., Melbourne-Thomas, J., Corney, S.P., Arrigo, K.R., Barbraud, C., Barnes, D.K., Bindoff, N.L., Boyd, P.W., Brandt, A., Costa, D.P., et al. (2014). Climate change and Southern Ocean ecosystems I: How changes in physical habitats directly affect marine biota. Glob. Change Biol. 20, 3004–3025 https://doi.org/10.1111/gcb.12623.

18. Crawshaw, L.I. (1979). Responses to rapid temperature change in vertebrate ectotherms. Am. Zool. 19, 225–237 https://doi.org/10.1093/icb/19.1.225.

19. Daniels, R.A. (1979). Nest guard replacement in the Antarctic fish *Harpagifer bispinis*: possible altruistic behaviour. Science 205, 831–833 https://doi.org/10.1126/science.205.4408.831.

20. Davison, W., Franklin, C.E. and Mckenzie, J.C. (1994). Hematological changes in an Antarctic teleost, *Trematomus bernacchii*, following stress. Polar Biol. 14, 463–466 https://doi.org/10.1007/BF00239050.

21. Domenici, P. (2010). Escape responses in fish: Kinematics, performance and behavior. In Fish locomotion: An eco-ethological perspective. (ed. P. Domenici and B.G. Kapoor, B.G.), pp. 123–170 Enfield, NH, USA, Science Publishers. https://doi.org/10.1201/b10190-6.

22. Domenici, P. and Hale, M.E. (2019). Escape responses of fish: a review of the diversity in motor control, kinematics and behaviour. J. Exp. Biol. 222, jeb166009 https://doi.org/10.1242/jeb.166009.

23. Domenici, P., Lefrançois, C. and Shingles, A. (2007). Hypoxia and the antipredator behaviours of fishes. Philos. Trans. R. Soc. B 362, 2105–2121 https://doi.org/10.1098/rstb.2007.2103.

24. Eastman, J.T. (1993). Antarctic fish biology: Evolution in a unique environment. San Diego, USA, Academic Press.

25. Eastman, J.T. (2017). Bathymetric distributions of notothenioid fishes. Polar Biol. 40, 2077–2099 https://doi.org/10.1007/s00300-017-2128-x.

26. Eastman, J.T. and Lannoo, M.J. (2004). Brain and sense organ anatomy and histology in hemoglobinless Antarctic icefishes (Perciformes: Notothenioidei: Channichthyidae). J. Morphol. 260, 117–140 https://doi.org/10.1002/jmor.10221.

27. Eaton, R.C., Bombardieri, R.A. and Meyer, D. (1977). The Mauthner-initiated startle response in teleost fish. J. Exp. Biol. 66, 65–81.

28. Eaton, R.C. and Emberley, D.S. (1991). How stimulus direction determines the trajectory of the Mauthner-initiated escape response in a teleost fish. J. Exp. Biol. 161, 469–487.

29. Eaton, R.C. and Hackett, J.T. (1984). The role of Mauthner cell in fast-starts involving escape in teleost fish. In Neural mechanisms of startle behavior (ed. R.C. Eaton), pp. 213–266. Boston, MA, USA, Springer https://doi.org/10.1007/978-1-4899-2286-1_8.

30. Eaton, R.C., Lee, R.K. and Foreman, M.B. (2001). The Mauthner cell and other identified neurons of the brainstem escape network of fish. Prog. Neurobiol. 63, 467–485 https://doi.org/10.1016/S0301-0082(00)00047-2.

31. Eliason, E.J. and Antilla, K. (2017). Temperature and the cardiovascular system. In Fish Physiology, Vol. 36B. The Cardiovascular System: Development, Plasticity and Physiological Responses (ed. A.K. Gamperl, T.E. Gillis, A.P. Farrell, and C.J. Brauner), pp. 235–297. San Diego, USA Academic Press, https://doi.org/10.1016/bs.fp.2017.09.003.

32. Fanta, E., Luchiari, P.H. and Bacila, M. (1989). The effect of temperature increase on the behavior of Antarctic fish. Proc. NIPR Symp. Polar Biol. 2, 123–130, http://id.nii.ac.jp/1291/00005059/.

33. Feller, G., Goessens, G., Gerday, C. and Bassleer, R. (1985). Heart structure and ventricular ultrastructure of hemoglobin- and myoglobin-free icefish *Channichthys rhinoceratus*. Cell Tissue Res. 242, 669–676 https://doi.org/10.1007/BF00225436.

34. Ferrando, S., Castellano, L., Gallus, L., Ghigliotti, L., Masini, M.A., Pisano, E. and Vacchi, M. (2014). Demonstration of nesting in two Antarctic icefish (*Genus Chionodraco*) using a fin dimorphism analysis and *Ex Situ* videos. PLoS ONE 9, e90512 https://doi.org/10.1371/journal.pone.0090512.

35. Franklin, C.E. and Johnston, I.A. (1997). Muscle power output during escape responses in an Antarctic fish. J. Exp. Biol. 200, 703–712.

36. Frederich, M. and Pörtner, H.-O. (2000). Oxygen limitation of thermal tolerance defined by cardiac and ventilatory performance in spider crab, *Maja squinado*. Am. J. Physiol. 279, R1531–R1538 https://doi.org/10.1152/ajpregu.2000.279.5.R1531.

37. Friedlander, M.J., Kotchabhakdi, N. and Prosser, C.L. (1976). Effects of cold and heat on behavior and cerebellar function in goldfish. J. Comp. Physiol. 112, 19–45 https://doi.org/10.1007/BF00612674.

38. Fry, F.E.J. and Hart, J.S. (1948). The relation of temperature to oxygen consumption in the goldfish. Biol. Bull. 94, 68–77 https://doi.org/10.2307/1538211.

39. Gollock, M.J., Currie, S., Petersen, L.H. and Gamperl, A.K. (2006). Cardiovascular and haematological responses of Atlantic cod (*Gadus morhua*) to acute temperature increase. J. Exp. Biol. 209, 2961–2970 https://doi.org/10.1242/jeb.02319.

40. Greenwood A.K., Peichel, C.L. and Zottoli, S.J. (2010). Distinct startle responses are associated with neuroanatomical differences in pufferfishes *J*. Exp. Biol. 213, 613–620. https://doi.org/10.1242/jeb.037085.

41. Grigaltchik, V.S., Ward, A.J.W. and Seebacher, F. (2012). Thermal acclimation of interactions: differential responses to temperature change alter predator-prey relationship. Proc. R. Soc. B 279, 4058–4064 https://doi.org/10.1098/rspb.2012.1277.

42. Hemmingsen, E.A. (1991). Respiratory and cardiovascular adaptations in hemoglobin-free fish: resolved and unresolved problems. In Biology of Antarctic Fish (ed. G. Di Prisco, B. Maresca, and B. Tota), pp. 191–203. Berlin, Germany, Springer https://doi.org/10.1007/978-3-642-76217-8_13.

43. Hancock, A. (1852). Observations on the nidification of *Gasterosteus aculeatus* and *Gasterosteus spinachia*. Ann. Mag. Nat. Hist. 10, 241–248 https://doi.org/10.1080/03745485609495690.

44. Holeton, G.F. (1970). Oxygen uptake and circulation by a hemoglobinless Antarctic fish (*Chaenocephalus aceratus, Lonnberg*) compared with three red-blooded Antarctic fish. Comp. Biochem. Physiol. 34, 457–471 https://doi.org/10.1016/0010-406X(70)90185-4.

45. Huey, R.B., Kearney, M.R., Krockenberger, A., Holtum, J.A., Jess, M. and Williams, S.E. (2012). Predicting organismal vulnerability to climate warming: roles of behaviour, physiology and adaptation. Philos. Trans. R. Soc. Lond. B 367, 1665–1679 https://doi.org/10.1098/rstb.2012.0005.

46. Hughes, G.M. and Roberts, J.L. (1970). A study of the effect of temperature changes on the respiratory pumps of the rainbow trout. J. Exp. Biol. 52, 177–192.

47. Heath, A.G. and Hughes, G.M. (1973). Cardiovascular and respiratory changes during heat stress in rainbow trout (*Salmo gairdneri*). J. Exp. Biol. 59, 323–338.

48. Hureau, J.C. (1985). Channichthyidae. In FAO species identification sheets for fishery purposes. Southern Ocean (Fishing Areas 48, 58 and 88). (CCAMLR Convention Area). Vol. 2, (ed. W. Fischer and J.C. Hureau) pp. 261–277. The Food and Agriculture Organization of the United Nations, Rome, Italy, available at http://www.fao.org/3/ah842e/ah842e00.htm.

49. Jayasundara, N., Healy, T.M. and Somero, G.N. (2013). Effects of temperature acclimation on cardiorespiratory performance of the Antarctic notothenioid *Trematomus bernacchii*. Polar Biol. 36, 1047–105 https://doi.org/10.1007/s00300-013-1327-3.

50. Joyce, W., Egginton, S., Farrell, A.P., Crockett, E.L., O’Brien, K.M. and Axelsson, M. (2018a). Exploring nature’s natural knockouts: in vivo cardiorespiratory performance of Antarctic fishes during acute warming. J. Exp. Biol. 221, pii: jeb183160 https://doi.org/10.1242/jeb.183160.

51. Joyce, W., Axelsson, M., Egginton, S., Farrell, A.P., Crockett, E.L. and O’Brien, K.M. (2018b). The effects of thermal acclimation on cardio-respiratory performance in an Antarctic fish (*Notothenia coriiceps*). Cons. Physiol. 6, coy069, https://doi.org/10.1093/conphys/coy069.

52. Jutfelt, F., Norin, T., Ern, R., Overgaard, J., Wang, T., McKenzie, D.J., Lefevre, S., Nilsson, G.E., Metcalfe, N.B., Hickey, A., et al. (2018). Oxygen- and capacity-limited thermal tolerance: blurring ecology and physiology. J. Exp. Biol. 221, pii: jeb169615 https://doi.org/10.1242/jeb.169615.

53. Kock, K. (1992) Antarctic fish and fisheries. Cambridge, UK, Cambridge University Press.

54. Liu, Y.C. and Hale, M.E. (2014). Alternative forms of axial startle behaviors in fishes. Zoology 117, 36–47 https://doi.org/10.1016/j.zool.2013.10.008.

55. Lucon-Xiccato, T., Nati, J.J., Blasco, F.R., Johansen, J.L., Steffensen, J.F. and Domenici, P. (2014). Severe hypoxia impairs lateralization in a marine teleost fish. J. Exp. Biol. 217, 4115– 4118 https://doi.org/10.1242/jeb.111229.

56. Mendonça, P.C. and Gamperl, A.K. (2010). The effects of acute changes in temperature and oxygen availability on cardiac performance in winter flounder (*Pseudopleuronectes americanus*). Comp. Biochem. Physiol. A 155, 245–252 https://doi.org/10.1016/j.cbpa.2009.11.006.

57. Meyers, J.R., Copanas, E.H. and Zottoli, S.J. (1998) Comparison of fast startle responses between two elongate bony fish with an anguilliform type of locomotion and the implications for the underlying neuronal basis of escape behavior. Brain Behav. Evol. 52, 7–22 https://doi.org/10.1159/000006548.

58. Nilsson, G.E., Rosén, P. and Johansson, D. (1993). Anoxic depression of spontaneous locomotor activity in crucian carp quantified by a computerized imaging technique. J. Exp. Biol. 180, 153–162.

59. O’Brien, K.M., Rix, A.S., Egginton, S., Farrell, A.P., Crockett, E.L., Schlauch, K., Woolsey, R., Hoffman, M. and Merriman, S. (2018). Cardiac mitochondrial metabolism may contribute to differences in thermal tolerance of red- and white-blooded Antarctic notothenioid fishes. J. Exp. Biol. 221, pii: jeb177816 https://doi.org/10.1242/jeb.177816.

60. Pacifici, M. Foden, W.B., Visconti, P., Watson, J.E.M., Butchart, S.H.M., Kovacs, K.M., Scheffers, B.R., Hole, D.G., Martin, T.G., Akcakaya, H.R., et al. (2015). Assessing species vulnerability to climate change. *Nat*. Clim. Change 5, 215–224 https://doi.org/10.1038/nclimate2448.

61. Pespeni, M.H. and Palumbi, S.R. (2013). Signals of selection in outlier loci in a widely dispersing species across an environmental mosaic. Mol. Ecol. 22, 3580–3597 https://doi.org/10.1111/mec.12337.

62. Pointer, M.A., Cheng, C.H., Bowmaker, J.K., Parry, J.W., Soto, N., Jeffery, G., Cowing, J.A. and Hunt, D.M. (2005). Adaptations to an extreme environment: retinal organisation and spectral properties of photoreceptors in Antarctic notothenioid fish. J. Exp. Biol. 208, 2363–2376 https://doi.org/10.1242/jeb.01647.

63. Pörtner, H.O. (2001). Climate change and temperature dependent biogeography: oxygen limitation of thermal tolerance in animals. Naturwissenschaften 88, 137–146 https://doi.org/10.1007/s001140100216.

64. Pörtner, H.-O., Bock, C. and Mark, F.C. (2017). Oxygen- and capacity-limited thermal tolerance: bridging ecology and physiology. J. Exp. Biol. 220, 2685–2696 https://doi.org/10.1242/jeb.134585.

65. Roche, D.G., Amcoff, M., Morgan, R., Sundin, J., Andreassen, A.H., Finnøen, M.H., Lawrence, M.J., Henderson, E., Norin, T., Speers-Roesch, B., Brown, C., Clark, T.D., Bshary, R., Leung, B., Jutfelt, F., Binning, S.A. (2020) Behavioural lateralization in a detour test is not repeatable in fishes. Anim. Behav. 167, 55–64; https://doi.org/10.1016/j.anbehav.2020.06.025

66. Rogers, L.J. (2010). Relevance of brain and behavioural lateralization to animal welfare. Appl. Anim. Behav. Sci. 127, 1–11 https://doi.org/10.1016/j.applanim.2010.06.008.

67. Ruud, J.T. (1954). Vertebrates without erythrocytes and blood pigment. Nature 173, 848–850 https://doi.org/10.1038/173848a0.

68. Sánchez-García, M.A., Zottoli, S.J. & Roberson, L.M. (2019). Hypoxia has a lasting effect on fast-startle behavior of the tropical fish *Haemulon plumieri*. Biol Bull. 237, 48–62 https://doi.org/10.1086/704337.

69. Schleidt, W.M. (1974). How “fixed” is the fixed action pattern? Z. Tierpsychol. 36, 184–211 https://doi.org/10.1111/j.1439-0310.1974.tb02131.x.

70. Schurmann, H. and Steffensen, J.F. (1994). Spontaneous swimming activity of Atlantic cod *Gadus morhua* exposed to graded hypoxia at three temperatures. J. Exp. Biol. 197, 129–142.

71. Sevenster, P. (1961). A causal analysis of a displacement activity (fanning in *Gasterosteus aculeatus*). Behaviour, (Suppl.) 9, 1–170 https://www.jstor.org/stable/30039143.

72. Shelford, V.E. and Powers, E.B. (1915). An experimental study of the movements of herring and other marine fishes. Biol. Bull. 28, 315–334 https://doi.org/10.2307/1536432.

73. Sillar, K.T. and Robertson, R.M. (2009). Thermal activation of escape swimming in post-hatching *Xenopus laevis* frog larvae. J. Exp. Biol. 212, 2356–2364 https://doi.org/10.1242/jeb.029892.

74. Somero, G.N. and DeVries, A.L. (1967). Temperature tolerance of some Antarctic fishes. Science 156, 257–258 https://doi.org/10.1126/science.156.3772.257.

75. Steinhausen, M.F., Sandblom, E., Eliason, E.J., Verhille, C. and Farrell, A.P. (2008). The effect of acute temperature increases on the cardiorespiratory performance of resting and swimming sockeye salmon (*Oncorhynchus nerka*). J. Exp. Biol. 211, 3915–3926 https://doi.org/10.1242/jeb.019281.

76. Tinbergen, N. (1951). The study of instinct. Oxford, UK, Clarendon Press.

77. Thomas, C.D., Cameron, A., Green, R.E., Bakkenes, M., Beaumont, L.J., Collingham, Y.C., Erasmus, B.F., De Siqueira, M.F., Grainger, A., Hannah, L., et al. (2004). Extinction risk from climate change. Nature 427, 145–148 https://doi.org/10.1038/nature02121.

78. Turner, J., Lu, H., White, I., King, J. C., Phillips, T., Hosking, J.S., Bracegirdle, T.J., Marshall, G.J., Mulvaney, R. and Deb, P. (2016). Absence of 21st century warming on Antarctic Peninsula consistent with natural variability. Nature 535, 411–415 https://doi.org/10.1038/nature18645.

79. Vallortigara, G. and Rogers, L.J. (2005). Survival with an asymmetrical brain: advantages and disadvantages of cerebral lateralization. Behav. Brain Sci. 28, 575–589 https://doi.org/10.1017/S0140525X05000105.

80. Van Iersel, J.J.A. (1953). An analysis of the parental behaviour of the male three-spined stickleback *(Gasterosteus Aculeatus L.) Behaviour*, Suppl., 3, 1–159 http://www.jstor.org/stable/30039128.

81. Vaughan, D.G., Marshall, G.J., Connolley, W.M., Parkinson, C., Mulvaney, R., Hodgson, D.A., King, J.C., Pudsey, C.J. and Turner, J. (2003). Recent rapid regional climate warming on the Antarctic Peninsula. Clim. Change 60, 243–274 https://doi.org/10.1023/A:1026021217991.

82. Ward, A.B. and Azizi, E. (2004) Convergent evolution of the head retraction escape response in elongate fishes and amphibians. Zoology (Jena). 107, 205–217 https://doi.org/10.1016/j.zool.2004.04.003.

83. Webb, P.W. (1976). The effect of size on the fast-start performance of rainbow trout *Salmo gairdneri*, and a consideration of piscivorous predator-prey interactions. J. Exp. Biol. 65, 157–177.

84. Webb, P.W. (1978). Fast-start performance and body form in seven species of teleost fish. J. Exp. Biol. 74, 211–226.

85. Wohlschlag, D.E. (1964). Respiratory metabolism and ecological characteristics of some fishes in McMurdo Sound, Antarctica. In Antarctic Research Series, *Vol.* 1. *Biology of the Antarctic Seas* (ed. M.O. Lee) American Geophysics Union, pp. 33–62 https://doi.org/10.1029/AR001p0033.

86. Zottoli, S.J. and Faber, D.S. (2000). The Mauthner cell: what has it taught us? The Neuroscientist 6, 25–37 https://doi.org/10.1177/107385840000600111

